# Regulatory network topology and the genetic architecture of gene expression

**DOI:** 10.1101/2025.08.12.669924

**Authors:** Matthew Aguirre, Jeffrey P. Spence, Guy Sella, Jonathan K. Pritchard

**Affiliations:** Department of Biomedical Data Science, Stanford University, Stanford CA; Department of Genetics, Stanford University, Stanford CA; Institute for Human Genetics, University of California, San Francisco, San Francisco CA; Department of Epidemiology & Biostatistics, University of California, San Francisco, San Francisco CA; Department of Biological Sciences, Columbia University, New York NY; Program for Mathematical Genomics, Columbia University, New York NY; Department of Biology, Stanford University, Stanford CA

## Abstract

In human populations, most of the genetic variance in gene expression can be attributed to *trans*-acting expression quantitative trait loci (eQTLs) spread across the genome. However, in practice it is difficult to discover these eQTLs, and their cumulative effects on gene expression and complex traits are yet to be fully understood. Here, we assess how properties of the genetic architecture of gene expression constrain the space of plausible gene regulatory networks. We describe a structured causal model of gene expression regulation and consider how it interacts with biologically relevant properties of the gene regulatory network to alter the genomic distribution of expression heritability. Under our model, we find that the genetic architecture of gene expression is shaped in large part by local network motifs and by hub regulators that shorten paths through the network and act as key sources of *trans*-acting variance. Further, simulated networks with an enrichment of motifs and hub regulators best recapitulate the distribution of *cis* and *trans* heritability of gene expression as measured in a recent twin study. Taken together, our results suggest that the architecture of gene expression is sparser and more pleiotropic across genes than would be suggested by naive models of regulatory networks, which has important implications for future studies of complex traits.

## 1 Introduction

Gene expression is widely thought to be an important mediator of the effects of non-coding genetic variation, which comprises the majority of hits from genome-wide association studies (GWAS) [1–3]. However, one major surprise from human genetic studies of gene expression has been that expression quantitative trait loci (eQTLs) close to a given gene tend to explain a minority of its genetic variance. This fraction of *cis*-acting heritability, 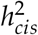, is estimated to have a genome-wide median of around 20% in studies of bulk tissue [4–7]. Similarly, *cis*-eQTLs explain a modest fraction of the heritability of many complex traits and diseases, and exhibit systematic differences with leading hits from genome-wide association studies (GWAS) [8, 9].

Still, *cis*-eQTLs have offered mechanistic insight into GWAS signals through approaches like statistical colocalization and transcriptome-wide association studies [10–13], even as it is debated whether most trait-relevant genetic variation can be discovered using assays of steady-state expression in bulk tissue [14]. But meanwhile, *trans*-eQTL discovery and analysis has been hampered by limited statistical power due to the higher multiple testing burden of genome-wide analysis and the likely smaller effect sizes of *trans*-eQTLs, compared to *cis*-eQTLs. Even though *trans*-acting variants are thought to explain the majority of gene expression heritability, their number and distribution throughout the genome remains unclear [6, 15].

At the same time, efforts to further map the genetic architecture of gene expression have seen growth in the size and resolution of functional genomics studies [3, 16–20], with concurrent development of inference methods that scale to these data [21–23]. These advances have increasingly relied on models of gene regulatory networks (GRNs) as more data are being used to assay real GRNs in many cell types and contexts. Specifically, network properties like modularity and regulatory hierarchy — respectively, the ideas that transcriptional master regulators control coherent gene sets and can direct cell-type differentiation — have been used to motivate considerations around cell-type composition effects in bulk and single-cell eQTL studies [16, 17, 22], as well as aggregation tests for *trans*-eQTLs [21, 24]. The relevance of these properties beyond gene expression has also been considered: in particular, genetic covariance induced by the structure of regulatory networks has been hypothesized to explain a substantial proportion of *trans*-acting heritability for complex traits [5, 6].

Here, we assess the implications of two simple and well-replicated observations about the distribution of expression heritability in bulk tissue — namely, that 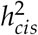 is low for the typical gene, and that *cis*-eQTLs have categorically larger effects than *trans*-eQTLs. We use a linear causal model of gene expression to build intuition about key local and global properties of regulatory networks. Next, we analyze the effects of these properties, showing how they can buffer or enhance the effects of *trans*-acting variation. Finally, we show that the observed distribution of *cis*-and *trans*-heritability constrains the space of plausible regulatory network structures, with the most realistic GRNs having a sparse architecture with master regulators and modular groups. In these GRNs, the bulk of *trans*-acting expression variance can be found along short paths in the network and at key genes with many pleiotropic effects. These features may therefore be useful in further mapping real genetic effects on gene expression.

## 2 Results

### 2.1 Genetic effects on gene expression

We motivate our work with data from a recent analysis of whole-blood gene expression from a twin study design to discover eQTLs and estimate heritability [7]. The data consist of eQTL summary statistics for 5,902 genes and heritability estimates for 11,409 protein coding genes measured in 1,497 individuals (**Methods**). Similar to other studies [4, 6, 16], the typical lead *cis*-eQTL effect size is 0.14 (standard deviations of expression), which is an order of magnitude greater than the typical lead *trans*-eQTL effect size (0.07; both are medians over 5,902 genes with a *cis*-eQTL, **Fig. 1A**). Meanwhile, the contributions to heritability are reversed, with *trans*-acting variation contributing the bulk of expression variance (**Fig. 1B**). The typical fraction of *cis*-acting heritability, 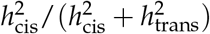, is 0.28 (median over 5,902 genes, **Fig. 1C**).

**Figure 1:**
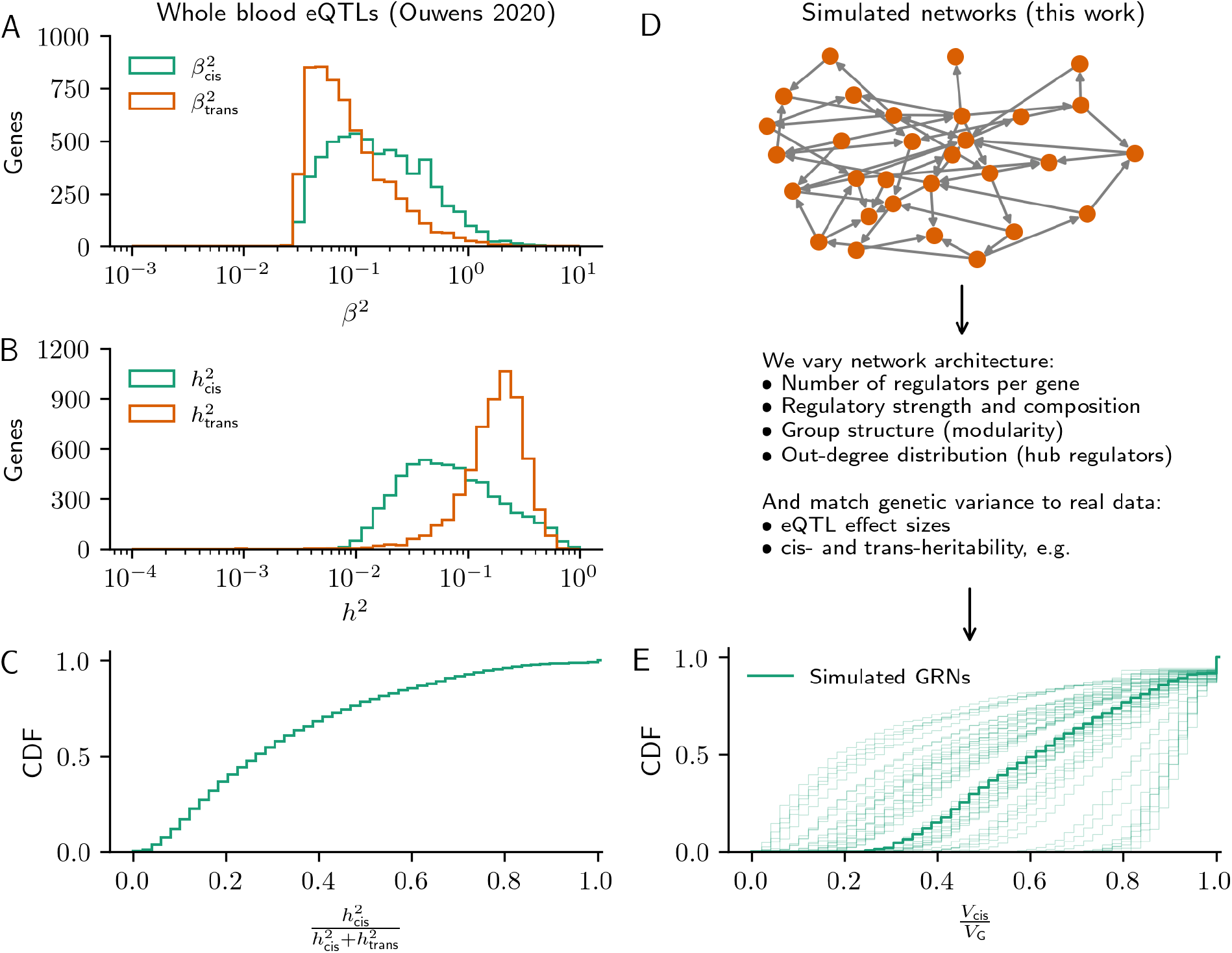
Overview of the study. **(A)** Lead eQTL effect sizes and **(B)** heritability from *cis*- and *trans*-acting genetic effects for 5,902 genes from a recent twin study of whole blood gene expression. **(C)** The cumulative distribution of *cis*-acting expression heritability from these same genes. **(D)**. Synthetic gene regulatory networks consist of a linear causal model on a directed acyclic graph, simulated using one of two generating algorithms (**Methods**). The parameters of these algorithms are standardized to tune interpretable properties of GRN architecture, and the expression model enables convenient computation of eQTL effect sizes and expression heritability. **(E)** Cumulative distribution of *cis*-acting expression heritability in 50 example synthetic GRNs (the overlay is the median distribution).

We use a two part model of how genetic variation affects gene expression through gene regulatory networks. The first component of the model is a graph generating algorithm, which we use to change the architecture of the causal network (of *trans*-acting regulators). Here, network architecture includes regulatory properties like the number and strength of regulators, and organizational properties like group structure and hierarchy (**Fig. 1D**; **Methods**). The second component of the model is a linear structural equation model of genetic effects on gene expression, which we use to measure the effects of network properties on the distribution of expression variance throughout the network. Specifically, we consider the size of typical lead *trans*-eQTL effects relative to lead *cis*-eQTL effects, 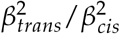, and the typical fraction of heritability due to *cis*-eQTLs, 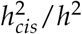 (**Fig. 1E**).

### 2.2 Local regulatory architecture

Under the assumption that genetic effects on gene expression are generally small, we make use a linear model of gene expression regulation. This model is a structural equation model (SEM) in which the expression, *y*, of a focal gene is determined by the effects of *q cis*-eQTLs (each with effect *β*_*i*_ from genotype *x*_*i*_) and *r* regulators (each with effect *γ*_*j*_):

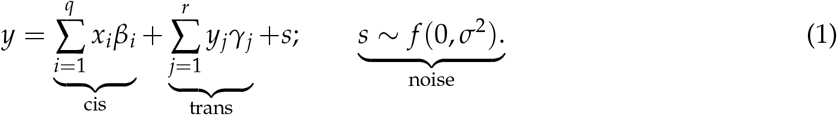

Here, for simplicity, we assume that all regulators have the same strength, *γ*, but that their effects can differ in sign: in particular, we assume gene *j* will act consistently as an activator for all of its targets with probability *p*^+^ and is otherwise a repressor. That is, with probability *p*^+^ the effect of a gene on the expression of all of its targets is +*γ*, and with probability (1 *− p*^+^) the effect is −*γ*. We further assume that *cis*-eQTLs are independent and, in total, contribute unit variance to each gene. The variance of the expression of the focal gene across individuals is then:

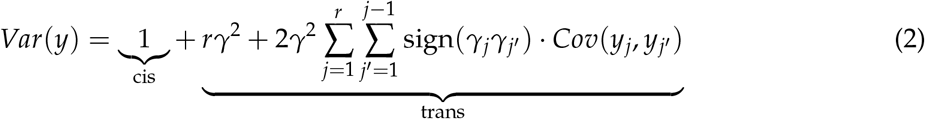

We will first use this model to show how local aspects of regulatory architecture — namely, the number, strength, and sign of regulators — affect how heritability and genetic effects on the expression of a focal gene are distributed among its regulators. Later, we will consider properties of entire regulatory networks. A more detailed description of the expression model and its assumptions can be found in **Methods**.

To show how properties of local regulatory architecture influence expression variance, we consider three different roles for one specific regulator of a gene, which we suppose has *r* + 1 regulators. If all of the regulators of this gene are independent, as in **Fig. 2A** (left), each regulator contributes *trans*-acting genetic variance with magnitude *γ*^2^, lowering the *cis*-fraction of heritability by the same amount (**Fig. 2B-C**). This is true regardless of how many of the regulators act as activators or repressors (**Fig. 2D, S2**).

**Figure 2:**
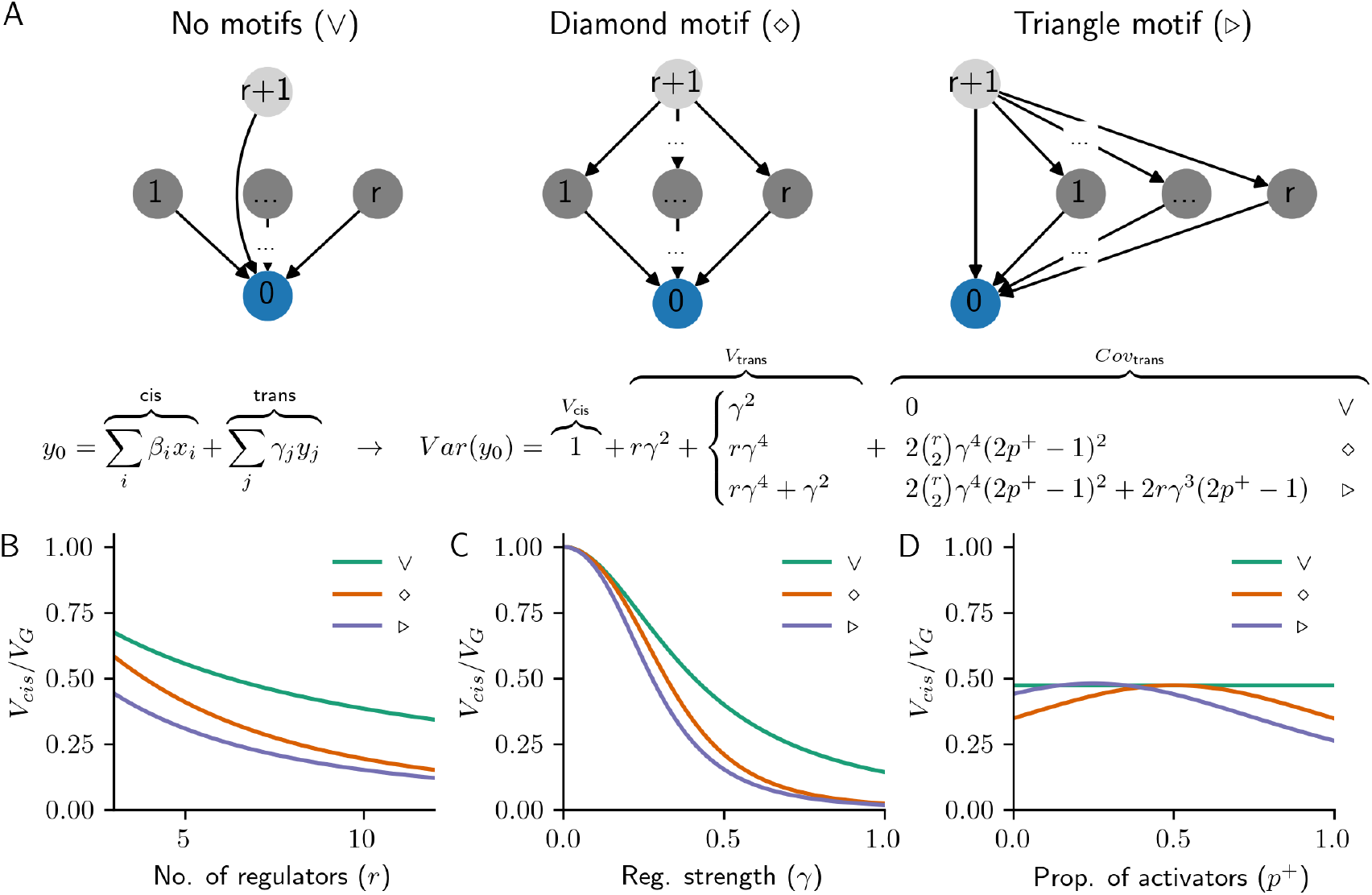
Local regulatory motifs. **(A)**. Linear causal model of *cis*- and *trans*-eQTL effects on a focal gene (0), with three different ways to add a new regulator (*r* + 1): as a direct regulator (no/V-motif), as a peripheral master regulator (diamond motif), or as both (triangle motif). We assume that *cis*-eQTLs contribute unit variance to each gene shown, which results in the *cis*- and *trans*-acting contributions to genetic variance given below the motif diagrams. Plots of this expression for each of the three motifs as a function of **(B)** the number of regulators per gene, *r*; **(C)** the strength of regulation, *γ*; and **(D)** the proportion of regulators that are activators, *p*^+^. Note that in the lower panels of the plot, unless otherwise stated, *r* = 6, *γ* = 0.4, and *p*^+^ = 1.

If the regulator is instead a peripheral master regulator (i.e., a gene that regulates the direct regulators of the focal gene), it can contribute *trans*-acting genetic covariance in addition to its direct effects on the focal gene (**Fig. 2A**, center and right). The expected magnitude and sign of this covariance depends not only on the number of existing regulators, *r*, and their strength, *γ*, but also on the fraction of activators, *p*^+^, and on whether the master regulator also directly regulates the focal gene (as in the “diamond” and “triangle” motifs; **Fig. 2B-D**; these are also respectively called “bi-parallel” and “feed-forward” motifs in the systems biology literature [25]). Further, these covariance terms are only nonzero in expectation over a random assignment of regulators as activators or repressors if these fractions are unequal (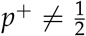; **Fig. 2D**; **Methods**).

Because of the covariance introduced by master regulators, it is possible for an indirect regulator of a gene to contribute more to its *trans*-acting genetic variance than one of its direct regulators (**Fig. 2B,C, S2**). In particular, if *p*^+^ is close to 1, motifs are more likely to be “coherent” (i.e., all paths from the master regulator to the target gene have the same sign [26]), and the expected *trans*-acting (co-)variance is larger (**Fig. S3**). Meanwhile, covariance introduced by “incoherent” motifs, where paths differ in sign [26], is expected to be negative. Hence, there is an important difference in how genetic effects flow through triangle and diamond motifs under activating and repressing regulation (since the chance of a motif being coherent or incoherent is related to powers of (2*p*^+^ *−* 1); **Fig. 2D**).

### 2.3 Modular network structure

Next, we consider the distribution of heritability and eQTL effects in the context of an entire GRN. Since we are interested in computing variances using a structural equation model, we simulate random directed acyclic graphs (DAGs) of causal regulatory relationships between genes. We do this using a standard random graph model, the planted partition model (PPM; **Methods**) [27]. Briefly, in the PPM, each of *n* nodes are assigned to one of *k* groups; edges exist between members of the same group with probability *p*, or different groups with probability *q*. To produce DAGs with this algorithm, nodes are randomly assigned indices, and edges are oriented to originate at the node with the lower index and point to the node with the higher index (**Methods**). Here, for ease in interpreting the structural properties of the network, we re-parameterize the model so that the expected number of regulators (edges) per gene is *r*, and the expected fraction of edges within groups is *m* (**Fig. 3A**,**S4, Methods**).

**Figure 3:**
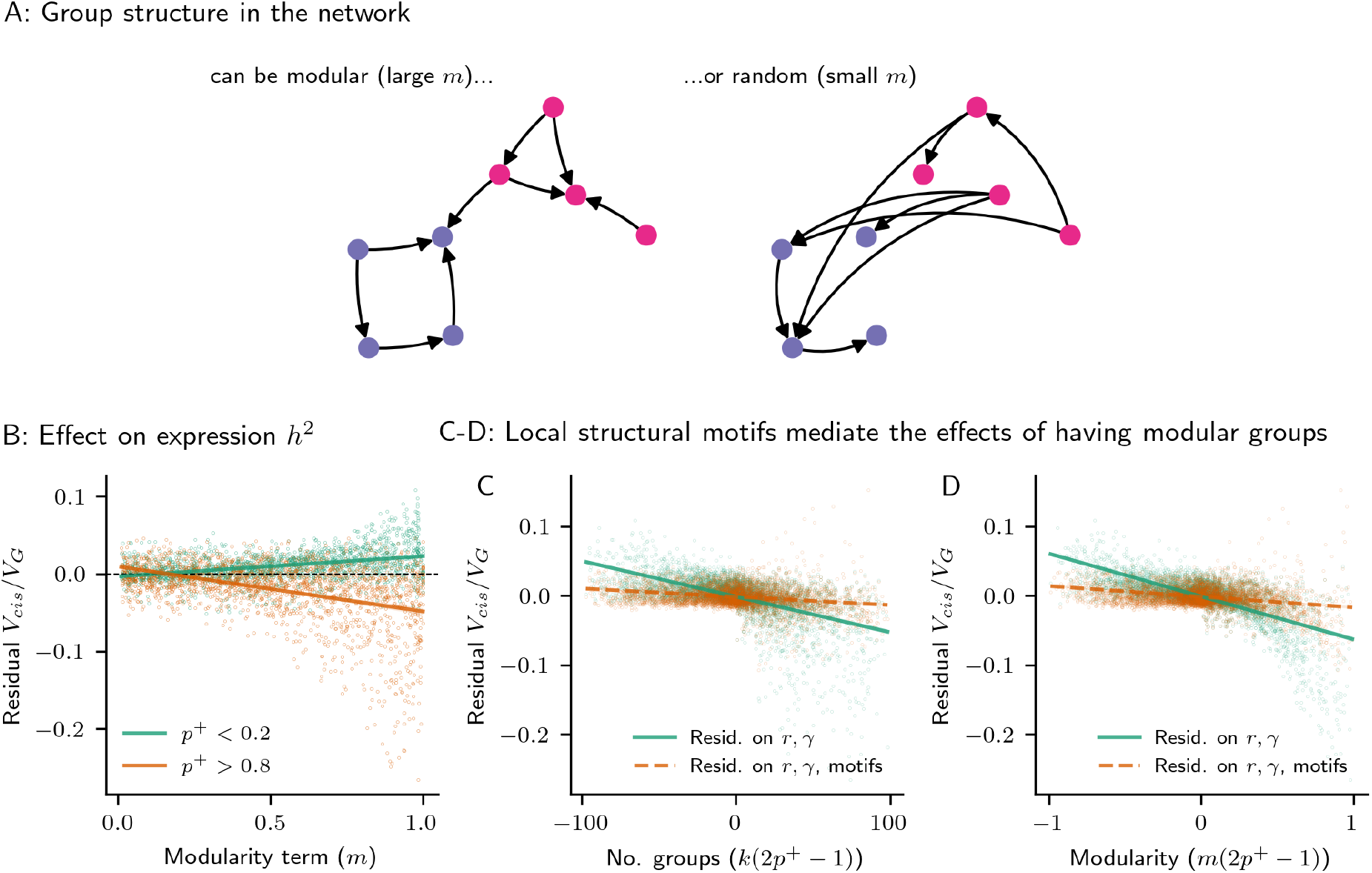
Modular network structure. **(A)**. Toy example of a GRN with modular groups that account for most edges in the network (large *m*), or groups that are random with respect to edges (small *m*). **(B)** The effect of modularity (*m*) on the median fraction of *cis*-acting genetic variance, residualized on direct effects from the number of regulators (*r*) and the strength of regulation (*γ*). The effect depends on the fraction of genes in the network that are activators, in a subset of 10,000 GRNs simulated using the planted partition model (PPM; **Methods**). **(C-D)** Structural motifs explain away the effect of the number of groups (*k*; **C**) and modularity (*m*; **D**), when appropriately scaled by the expected sign of regulation (2*p*^+^− 1) — residualizing the median fraction of *cis*-acting genetic variance in the GRN by the number of triangle and square motifs nearly eliminates the relationship with the scaled group structure terms.

We use the relationships specified by the simulated DAG as the basis for the gene expression model. As in the previous section, the expression, *y*_*g*_, of gene *g* is

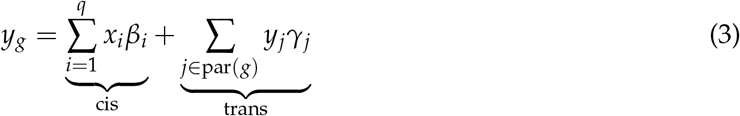

where par(*g*) denotes the “parents” of gene *g* — nodes with edges that point into node *g*. We again assume that *γ*_*j*_ has fixed magnitude *γ*, that each gene acts consistently as an activator with probability *p*^+^ or as a repressor with probability (1− *p*^+^), and that all genes in the GRN have *cis*-eQTLs that collectively contribute unit variance to their expression.

To describe the effects of local and global network properties on expression heritability, we simulated 10,000 synthetic DAGs using the PPM and computed the resulting fraction of *cis*-heritability, 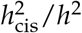, for every gene in the network (**Methods**). We used the median of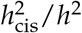 across genes as a summary statistic for each GRN, and considered how this median *cis*-heritability fraction changes as a function of the parameters of the PPM and the gene expression model.

As was the case for a single gene, strong local regulation reduced the fraction of *cis*-acting heritability across networks. This was true both for the number of regulators *r* and their strength *γ* — in particular, 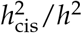 was tightly related to 1/(1 + *rγ*^2^) (**Fig. S5**). After regressing out this direct effect, the other parameters had more obvious effects: the fraction of genes that are activators, *p*^+^, increased contributions from *trans*-eQTLs and scaled the contribution of triangle and diamond motifs to *trans*-acting variance (**Fig. S5**).

Further, group structure also had an important effect on the magnitude of indirect *trans*-acting genetic variance. Similar to motifs, the number of groups, *k*, and their modularity, *m*, interacted with the fraction of activators, *p*^+^, to explain some residual variance in *cis*-heritability fraction. In networks with many activators (*p*^+^ near 1), modularity amplified *trans* effects, but in networks with many repressors, modularity dampened them (**Fig. 3B**). Since *m* is strongly related to the number of triangle and diamond motifs in the network (**Fig. S6**), we conducted a mediation analysis to consider the extent to which these motifs explain away the effects of group structure. In these networks, nearly all of the statistical variance explained by group structure (*k* and *m*, scaled by (2*p*^+^ − 1)) is explained away by triangle and diamond motifs (**Fig. 3C,D**; model *R*^2^ = 0.177, 0.260 without regressing out motifs, and model *R*^2^ = 0.010, 0.020 when regressing out motifs, respectively for groups and modularity). Thus, modular groupings altered global patterns of expression heritability in the network by introducing local structures.

### 2.4 Regulatory hubs

In biological networks across species, regulatory activity is thought to concentrate at hubs with downstream effects on modular groups [16, 28]. We therefore expect GRNs to have heavy-tailed out-degree distributions. To assess how hubs affect *cis* and *trans* heritability, we modified a previous directed scale-free network generating algorithm to produce acyclic graphs (**Methods**) [29,30]. The parameters of this algorithm are identical in interpretation to the PPM, but with an additional out-degree uniformity term, *d*. Smaller values of *d* produce networks with a heavy-tailed out-degree distribution, while larger values of *d* produce networks with a less hub-like regulatory architecture (**Fig. 4A, S7**).

**Figure 4:**
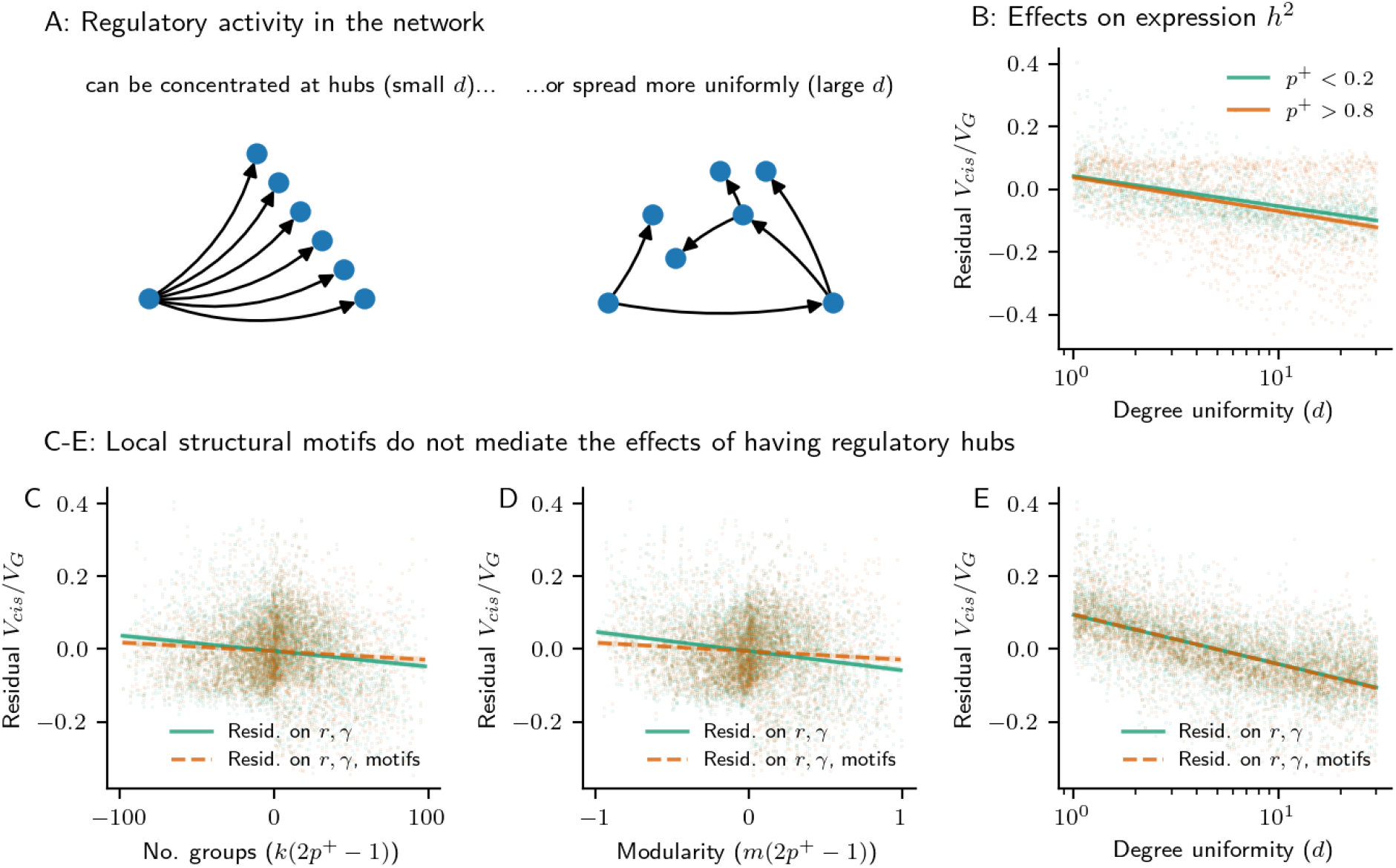
Regulatory hubs in the GRN. **(A)**. Toy example of a GRN with hub regulators (small *d*) or with a more uniformly spread regulatory architecture (large *d*) **(B)** The effect of degree uniformity, *d*), on the median fraction of *cis*-acting genetic variance, residualized on direct effects from the number of regulators (*r*) and the strength of regulation (*γ*). The effect no longer on the fraction of genes in the network that are activators, in a subset of 10,000 GRNs simulated using our generating algorithm (**Methods**). **(C-E)** Structural motifs explain away the effect of **(C)** the number of groups, *k*, and **(D)** modularity, *m*, when appropriately scaled by the expected sign of regulation (2*p*^+^− 1), but not **(E)** the effect of degree uniformity, *d*. Further residualizing the median fraction of *cis*-acting genetic variance in the GRN by the number of triangle and square motifs nearly eliminates the relationship with the scaled group structure terms, but has no effect on the relationship with degree uniformity.

In networks generated using this model, the effects of both local and global regulatory properties (number and strength of regulators, number of triangle and diamond motifs, and modular group structures) were consistent with results from the PPM model. Namely, direct regulatory effects had the largest impact on the *cis*-acting fraction of heritability (in particular 1/(1 + *rγ*^2^), which we again regress out for downstream analysis; **Fig. S8**). Furthermore, the fraction of genes that are activators, *p*^+^, was still a significant driver of *trans*-acting heritability and retained significant interactions with other structural properties of the GRN (**Fig. S8**).

We also found that out-degree dispersion reduced the *cis*-fraction of expression heritability in a manner which does not depend on *p*^+^ (**Fig. 4B**). Further, this effect was not mediated by the number of motifs in the network (**Fig. 4E**), although motifs were important mediators of the effects of modular groups (**Fig. 4C,D**). Rather, hub-like regulatory architecture decreased *trans*-acting heritability even as these networks were motif-rich compared to networks with a dispersed regulatory architecture (unless the network was dense and had many modular groups, **Fig. S9**). Instead, introducing hubs into the architecture of the network (i.e., lowering *d*) resulted in fewer long paths through the GRN, which lowered the number of (total) regulators per gene and meant that regulators closer to genes explained more *trans*-heritability (**Fig. S10**).

### 2.5 Network distribution of heritability

We have so far considered how network properties alter the distribution of expression heritability for the median gene in the GRN. But this raises a key question: which of these properties, if any, are necessary to explain the observed distribution of *cis*- and *trans*-heritability?

To address this question, we returned to our motivating dataset [7]. We compared synthetic networks generated using the PPM and scale-free network models to the observed distribution of *cis*-heritability fraction across genes in these data, and evaluated characteristics of the 250 GRNs which are closest to it (K-S test; **Methods**).

Overall, we found that the observed distribution of *cis*-heritability was well-matched by some networks generated by the scale-free network generating algorithm (**Fig. 5A**), while networks from the PPM tended to be less similar (**Fig. S11**). Since we did not assume a particular distribution of *cis*-eQTL effect sizes for the synthetic GRNs, we compared the contribution of the largest *trans*-regulator with the *cis*-acting variance for each gene as a proxy for *cis*- and *trans*-eQTL effect sizes. This corresponds to the assumption that lead *cis*-eQTL effects are similarly distributed for all genes and that all *trans*-eQTLs are also *cis*-eQTLs. Even without explicitly matching simulated networks on the distribution of *trans*-eQTL effect sizes, we found that the simulated GRNs that were well-matched to the observed distribution of *cis*-heritability fraction also tended to match the observed ratio of median lead *trans*-to *cis*-eQTL effect sizes (median 0.51 in matched GRNs; **Fig. 5B**).

**Figure 5:**
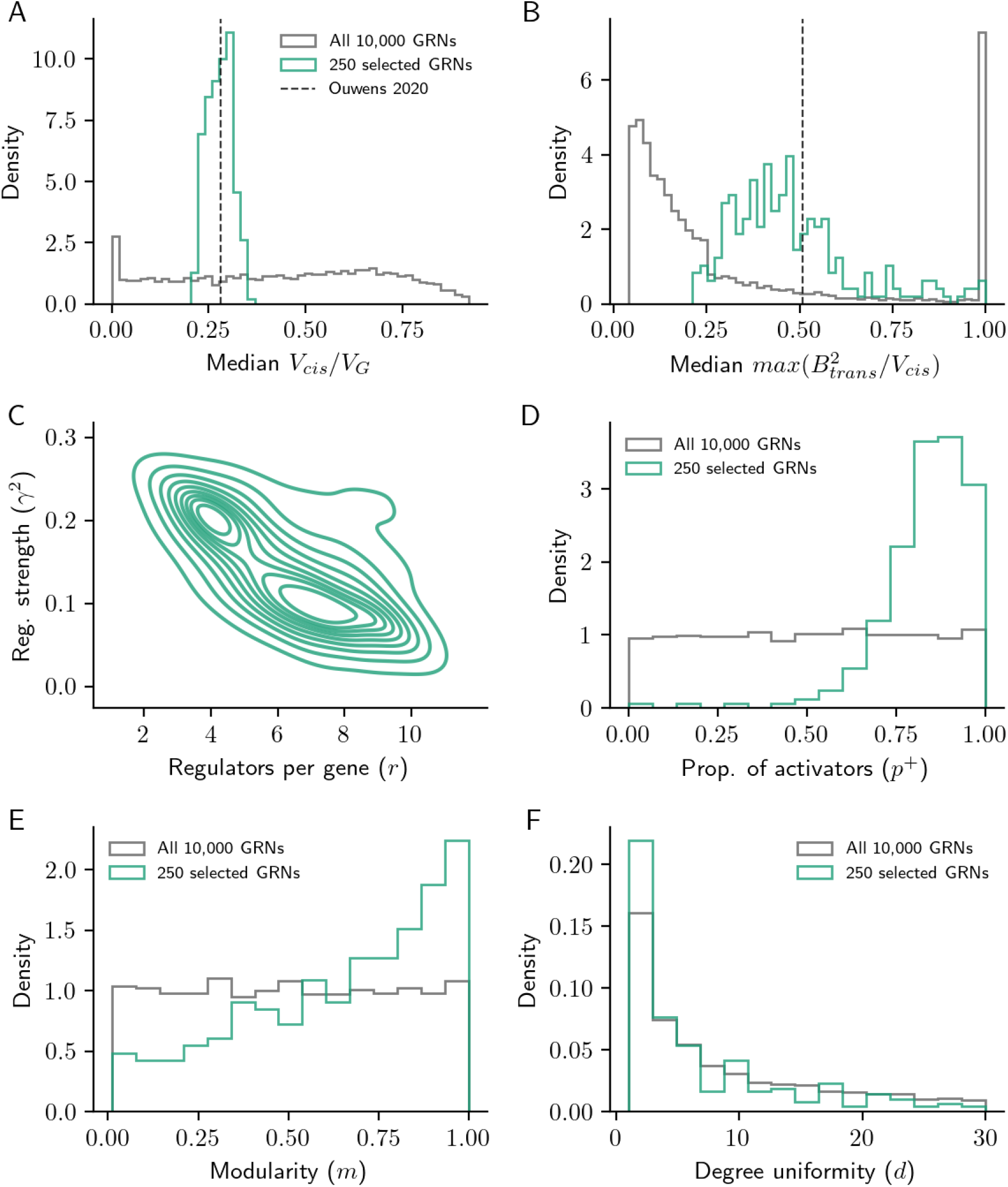
Relationship to real data. Median fraction of **(A)** *cis*-acting expression variance and **(B)** scaled *trans*-acting effects in all 10,000 scale-free GRNs, and in the 250 GRNs most closely matched to the cumulative distribution of 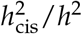 from real data. Distribution of **(C)** local regulatory parameters (*r* and *γ*); **(D)** fraction of activators (*p*^+^); and structural parameters for **(E)** modularity (*m*); and **(F)** degree uniformity (*d*) in the matched networks, against the background distribution of GRN parameters.

Further, we found that the GRNs matched to real data share characteristic properties. First, while *r* and *γ* vary in the set of well-matched networks, the quantity 1/(1 + *rγ*^2^) is tightly constrained, indicating that the distribution of *cis*-acting heritability is highly informative about the strength of direct regulation (**Fig. 5C**). Similarly, the fraction of activators *p*^+^ tends to be at the high end of its range, mirroring the empirical observation that activation is a more common regulatory mechanism than repression (**Fig. 5D**) [26]. Further, when *p*^+^ is large in our model, motifs like the feedforward loop are more likely to be coherent than incoherent. This would run counter to the view that incoherent feedforward loops are key motifs in biological networks [26], but it may be instead that these motifs are structured (rather than random) with respect to the sign of each regulator in real GRNs.

Finally, the well-matched GRNs tend to be modular and have hub regulators, as evidenced by a minor enrichment of *m* near 1 and *d* at the low end of its range (**Fig. 5E,F**). These parameters, as with many terms in our model, have largely independent effects on how well GRNs match real data (**Fig. S13**). However, after adjusting for the effects of the modularity and degree parameters (*m,d*), we find only weak statistical evidence for an enrichment of structural motifs among matched networks (triangles and diamonds; **Fig. S12**).

In previous work, we showed that experimental perturbation data also suggested that GRNs are sparse, modular, and have hub regulators [30]. On balance, the effects of these parameters in this setting was to dampen the effects of a random perturbation on the expression state of the network. Here, we find that these same properties are also associated with weaker *trans*-eQTL effects in aggregate (as measured by the fraction of *cis*-acting heritability; **Fig. S12**). Thus these two distinct modeling frameworks, matched to two distinct types of data, both indicate that sparsity, modularity, and hub regulators are important structural properties of GRNs that coherently affect key measures of their function.

### 2.6 Implications for discovery

Finally, we consider the implications of these characteristic properties of GRNs for the genetic architecture of gene expression. For this, we compared how expression heritability is distributed throughout the network in three exemplar GRNs that have diverse structural features — one GRN with realistic scale-free structure and which closely matches summaries of the twin study estimates of expression heritability; another GRN with similar properties to this network, but lacking regulatory hubs (large *d*); and a third GRN lacking modular structure, but with a similar number of edges (PPM). Although these networks have a similar median 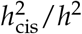, their distributions of *cis* heritability fraction are otherwise quite different and only the first GRN matched real data (**Fig. 6A**).

**Figure 6:**
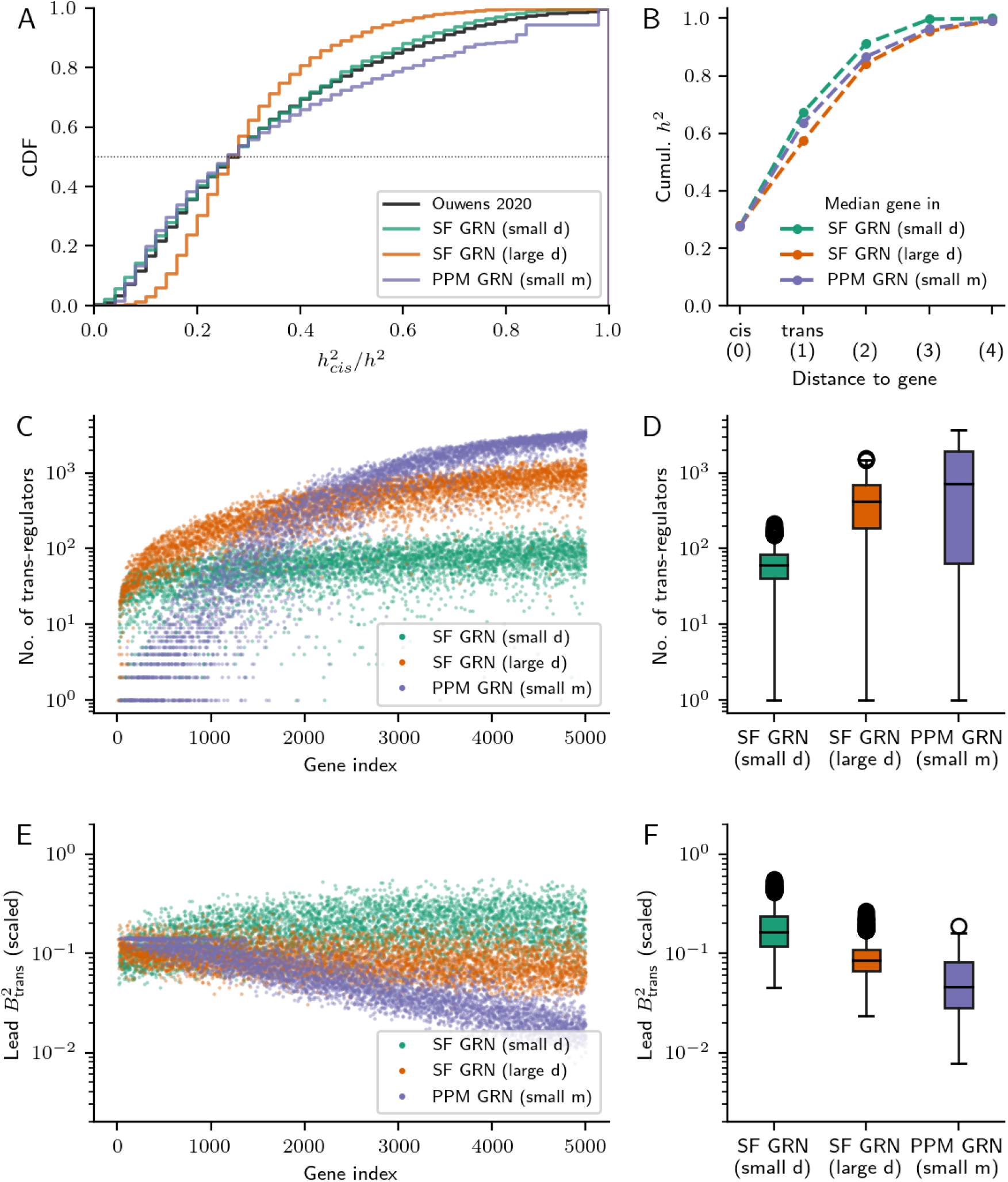
Genetic architecture in diverse GRNs. **(A)**. Cumulative distribution of the fraction of *cis*-acting expression variance in three example GRNs with different structural properties, against the distribution from real data (Ouwens *et. al*., 2020). **(B)** Median cumulative heritability as a function of network distance in three example GRNs. Median is over genes in the network; *cis*-effects are at distance 0, *trans*-effects from direct regulators are at distance 1, etc. **(C)** Number of *trans*-regulators (direct and indirect) and **(E)** lead *trans*-eQTL effect sizes for all genes in the example GRNs, as a function of a gene’s position (“index”) in the topological ordering of the underlying DAG. **(D**,**F)** Histograms of the distribution in (C,E).

These GRNs were further differentiated when considering contributions to heritability beyond *cis* and *trans*. For this, we decomposed expression variance as a function of distance in the GRN to a focal gene. In the well-matched scale-free GRN, 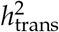 was modestly more attributable to regulators closer to their target genes in the GRN (i.e., at distance 1 or 2 than distance 3 or 4; **Fig. 6B, S14, S15**). Regulators within two hops of a focal gene (e.g., those as diagrammed in **Fig. 2**) cumulatively explained 91.2% of heritability for the median gene in the well-matched GRN, compared to 84.3% or 86.6% in the GRNs without hubs or modular groups.

This concentration of expression heritability near a given gene in the well-matched GRN occurred despite the typical gene in it having fewer regulators (**Fig. 6C-D**). Though genes in all three GRNs were expected to have a similar number of direct regulators, (*r* = 7.93,7.93, and 8.04 across the three GRNs), their varied structural properties dramatically altered the architecture of indirect regulation. Genes near the top of the PPM DAG had tens (or fewer) of upstream regulators, but genes near the bottom sometimes had thousands (**Fig. 6C**) — meanwhile, the number of regulators of genes in both scale-free GRNs tended to plateau towards the bottom of the DAG (**Fig. 6C**). Similarly, a gene in the scale-free GRN with hubs typically had substantially fewer upstream regulators (median 61) than genes in either other GRN (median 413 in the scale-free GRN without hubs, 589 in the PPM GRN; **Fig. 6D**). We further measured this difference in indirect regulatory architecture by decomposing heritability as a function of distance between regulators and target genes in each GRN; the fraction of variance explained by regulators at various distances also stabilized towards the bottom of the DAG in the scale-free GRNs, but not the PPM GRN (**Fig. S14**).

These exemplar GRNs also had different distributions of lead *trans*-eQTL effect sizes across genes in the network (**Fig. 6E-F**). As in real eQTL studies, we computed these effects in units of expression standard deviation, separately for each gene in the network. We found that across networks, genes with more upstream regulators typically had smaller (scaled) leading *trans*-eQTL effects, since each regulator contributed a smaller proportion of variance. This was particularly true for genes near the bottom of the PPM GRN (**Fig. 6C,E**). Meanwhile, in the comparatively motif-rich scale-free networks, where the number of upstream regulators grew more slowly as a function of gene index (**Fig. 6C**), the distribution of *trans*-eQTL effect sizes was more uniform across genes (**Fig. 6E,F**).

Overall, lead *trans*-eQTL effects were largest in the GRN that was well-matched to real data, and smaller in the matched GRNs that had a similar median 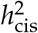 but lacked hub regulators and modular groups (**Fig. 6F**). We found this to be somewhat counterintuitive, as low out-degree uniformity (small *d*) and high modularity (large *m*) were marginally associated with lower *trans*-acting variance (i.e., with higher median 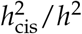; **Fig. S8**). However, the median 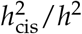 of genes in a GRN does not specify the full distribution of 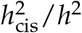 (**Fig. 6A**), nor does it specify the underlying architecture of *trans*-effects (**Fig. 6F**). This distribution of *trans*-eQTL effect sizes depends on the number and nature of paths between a gene’s upstream regulators, which is dramatically altered by modular group structure or hub regulatory architecture. Taken together, these network properties suggest that *trans*-acting effects could be larger and closer to focal genes than would be naively expected under an unstructured GRN model with low *cis*-heritability, with fewer loci contributing *trans*-acting variance.

## 3 Discussion

In this work, we have characterized the space of gene regulatory networks that are compatible with a simple observation about the genetic architecture of gene expression: namely, that 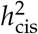 is generally low despite 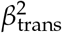 usually being small. To do this, we evaluated the effects of various structural properties of GRNs, and their necessity (or lack thereof) to match patterns in real data. Our results illuminate what eQTLs can tell us about incompletely mapped GRNs, and what this space of plausible GRNs tells us about the eQTLs we are yet to discover.

Across networks, we found that direct regulatory effects (i.e., the number of regulators per gene and the average strength of regulation) have a key role in determining the relative sizes of *trans*-eQTL effects and the proportion of *trans*-acting heritability. However, peripheral effects due to the organization of distal regulators were also important for matching the observed genetic architecture of genome-wide gene expression. In particular, we found that local motifs mediate a substantial portion of the effects of group structure in the network, and that regulatory hubs further located *trans*-eQTL effects closer to genes and at key loci in the network.

While our modeling captures the key features of GRNs, it does make several simplifying assumptions about the functional form of gene expression and the activity of regulators throughout the GRN. In reality, gene expression is a dynamic, time-varying process with non-linearities due to molecular kinematics and protein interactions; regulatory effects are not all equal in magnitude, and regulators do not necessarily act with consistent sign, nor are local motifs random with respect to this direction of effect; and finally, not all genes have equal *cis*-acting variance, and the distribution of *cis*-eQTL effects is of further interest in the context of complex trait genetic architecture. While many of these parameters are also often abstracted away in quantitative genetics models (e.g., the infinitesimal model is agnostic to regulatory architecture), they are important aspects of GRNs and their function across time and tissues. Relatedly, our work does not resolve questions about eQTLs whose effects may only be visible over dynamic or developmental trajectories, or in narrower populations of single cells in bulk tissues. The question of why some genes lack *cis*-eQTL variance in certain contexts also remains open for further study.

Still, it remains that the structural properties of GRNs exert significant influence on the distribution of genetic effects on gene expression. In particular, synthetic GRNs with structural features characteristic of biological networks (i.e., sparsity, modularity, and a heavy-tailed out-degree distribution) were categorically better matched to real data than GRNs without them. The effects of these features on the distribution of heritability through a GRN also suggest that the architecture of gene expression is sparser than would be anticipated under a unstructured model of GRN properties. Recent work using genome-scale Perturb-seq has also found that most regulators influence the expression of a small number of downstream genes in the cell (45 for the median gene, 500 or more for the typical essential gene [31]). This finding is consistent with our result that there are tens of regulators upstream of the median gene in a synthetic GRN with biological structural features and a realistic distribution of expression heritability (**Fig. 6D**).

We likewise found that *trans*-eQTLs should be more concentrated and have more clustered downstream effects than would be suggested by unstructured graph models. Our results therefore support the use of statistical aggregation tests that exploit these features of GRNs to further map *trans*-acting genetic effects on gene expression. To the extent that this co-regulation reflects functional relatedness, we further anticipate that using this genetic signal to define gene “programs” will be a key way forward to uncover the biology of complex traits and diseases.

## Acknowledgements

We would like to thank members of the Pritchard Lab at Stanford University for helpful comments and discussion related to this work. M.A. acknowledges support from a Microsoft Research PhD Fellowship, and from the National Library of Medicine (NLM) under training grant T15LM007033. This work was supported by the National Human Genome Research Institute (NHGRI) under grants R01HG008140, R01HG014005, and U01HG012069 (J.K.P.) and by the National Institute of General Medical Sciences (NIGMS) under grant R01GM115889 (G.S.).

## 4 Methods

### 4.1 Whole-blood eQTL data

In this work, we make use of data from a study of whole blood gene expression in 1,497 individuals from 709 twin pairs (459 monozygotic and 150 dizygotic) [7]. Study design and data collection procedures are described in the initial publication. The data consist of measurements of 52,844 genes, which were subsequently filtered for being protein coding, having read counts above zero in at least 85% of samples in each zygosity group (i.e., expressed in in at least 780 MZ twins and 255 DZ twins), a median expression count above 10, and more than 20 SNPs in the cis-window, resulting in an analysis of 11,409 genes. Heritability was estimated using the software tool GCTA [32], with estimates partitioned into *cis*- and *trans*-acting components by considering variation within a 250 kilobase window flanking the gene body as acting in *cis* (and in *trans* otherwise). These data were used in the format provided in Supplementary Table 2 of the initial publication [7].

We computed the *cis*-fraction of expression heritability by dividing the point estimates of *cis*- and *trans*-heritability from GCTA as 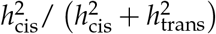, and further subsetted the data to the 5,092 genes with at least one *cis*- and *trans*-eQTL called in the study. This additional filtration step removes 1,293 genes with estimates of 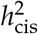 or 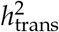 which were indistinguishable from zero, and introduces an upward shift in the distribution of *cis*-heritability (but not *trans*-heritability; **Figure S1**).

### 4.2 Gene expression model

We model gene expression using a linear structural equation model using a set of causal gene regulatory relationships specified by a directed acyclic graph (DAG) between *n* genes. We consider the nature of the DAG and its structural properties in the ensuing sections — to start, we assume that gene *i* harbors *q*_*i*_ *cis*-acting quantitative trait loci (cis-eQTLs; *q*_*i*_ *>* 0) and is influenced by some number of regulators, *r*_*i*_ (*r*_*i*_ *≥* 0). Denoting the expression of gene *i* in a randomly chosen individual by *y*_*i*_, we have that

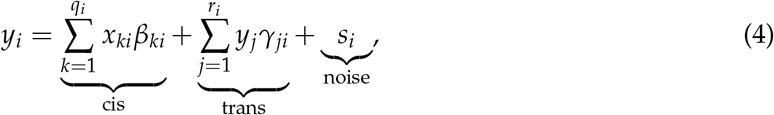

where *β*_*ki*_ and *x*_*ki*_ are respectively the effect of the *k*^th^ eQTL for gene *i* and an indicator variable for its genotype; *γ*_*ji*_ and *y*_*j*_ are the effect of the *j*^th^ regulator of gene *i* and its expression respectively; and *s*_*i*_ is non-genetic noise with zero mean and variance 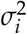, assumed to be independent of genotypes and uncorrelated across genes.

We further assume that each regulator acts consistently as an activator or repressor and has the same magnitude of effect (i.e., *γ*_*ji*_ = *p*_*j*_*γ*, with *p*_*j*_ denoting the sign for gene *j*); and that the variance due to *cis*-acting eQTLs and non-genetic noise in expression are the same for all genes (respectively 1, and *σ*^2^). The variance for gene *i* is then

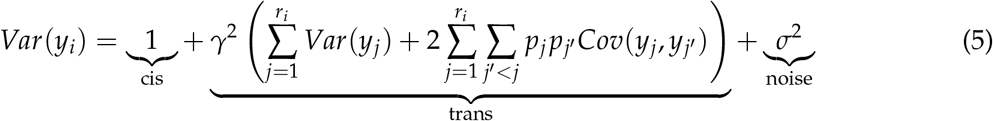

With these assumptions, we see that *γ*^2^ scales the relative contributions of *cis*- and *trans*-acting genetic variation, and *σ*^2^ scales the ratio of genetic and non-genetic variance (i.e., the heritability). Specifically, we have that *h*^2^ = 1/(1 + *σ*^2^) for all genes in the network, since the *trans*-acting terms above can be expanded in terms of *cis*-acting terms for other genes (i.e., the ratio of incoming genetic and noise terms is the same). We can then write that

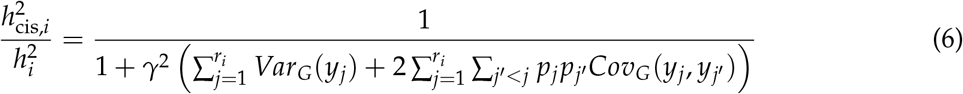

with *Var*_*G*_ and *Cov*_*G*_ respectively denoting the genetic variance and genetic covariances of genes *j* and *j′*. A complete derivation is in **Appendix A**.

For the example motifs in **Fig. 2**, we can further take the expectation over the random assignments of genes as activators or repressors (i.e., the signs of the edge weights). In this setting, we have the following equations for the expected *cis*-contribution to variance over the expected total genetic variance:

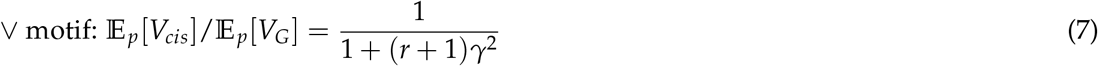

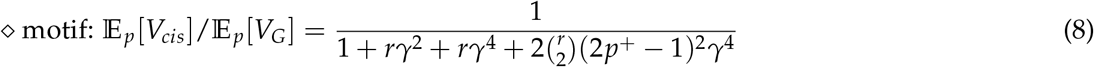

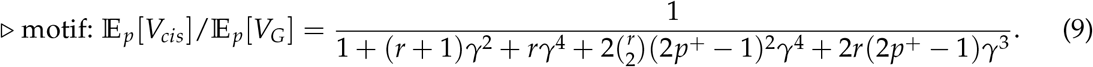

where *p*^+^ is the probability that each gene is an activator as opposed to a repressor.

### 4.3 Planted partition model

We use the planted partition model (PPM) to produce the synthetic directed acyclic graphs (DAGs) which form the basis of our simpler GRN model. The PPM is a simple extension of the binomial (Erdos-Renyi /ER) random graph model to include group structure [27, 33]. The ER model has two parameters: the number of nodes, *n*, and the edge probability, *p*. In this model, all edges are independent and identically distributed Bernoulli random variables with parameter *p*. The PPM extends this by assigning the *n* nodes to one of *k* groups uniformly at random; edges exist between members of the same group with probability *p* and between members of different groups with probability *q*.

To produce DAGs using the PPM, we further generate a random topological ordering of the *n* nodes. Edges are directed from the node earlier in the ordering to the node later in the ordering; this constraint also prevents the creation of cycles.

For consistency in interpreting parameters of the gene expression model, and to account for imposition of acyclicity, we reparameterize the PPM in terms of *r, k*, and *m* — respectively, the expected number of regulators per gene, the number of groups in the network, and the expected fraction of edges drawn between nodes belonging to the same group. In terms of the original parameters *p* and *q*, we have

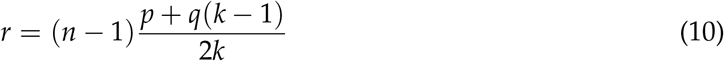

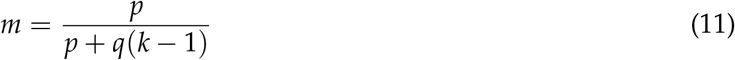

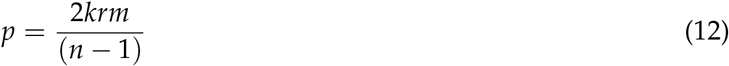

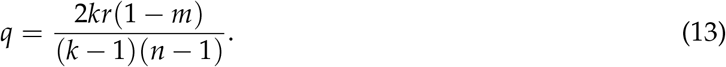

We note that there is degeneracy in the above if *k* = 1; in this case, we have recovered the ER model. Further, in the typical PPM formulation, *p < q* means that the network groups are *dissociative* (i.e., nodes tend to be connected with members of other groups rather than their own); with the above parameters, this occurs when *m <* 1/*k*. This case is not of interest here, and so we enforce *m >* 1/*k* for simulations.

The output of the PPM is a DAG with *n* genes assigned into *k* groups. With the above parameters, the groups account for roughly *m* percent of the edges in the network, and the typical gene has *r* regulators (**Fig. S4**). To simulate gene expression from this network, we fix the strength of regulation at *γ* and independently assign each gene a regulatory sign, which is positive with probability *p*^+^. Taking *G* to be the weighted adjacency matrix after this process, we have that the fraction of *cis*-acting genetic variance for the *i*’th gene in the GRN is

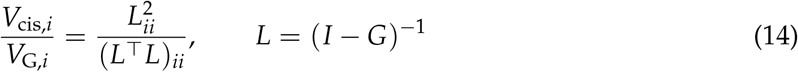

where (*X*)_*ii*_ is the *i*^th^ entry on the diagonal of the matrix *X* and *I* is the *n*-dimensional identity matrix. A complete derivation of this result is in **Appendix A**.

#### 4.3.1 Choosing hyperparameters

To produce the example distribution of heritability in **Fig. 1**, we simulated 50 GRNs with DAG structures produced by the PPM using generating parameters as given below:

- Number of genes *n* = 1000
- Number of groups *k* = 1
- Modularity term *m* is not applicable in this case (these are ER graphs)
- Expected regulators per gene *r ∼* Uniform(4, 8)
- Strength of regulation *γ ∼* Uniform(0.1, 0.5)
- Expected fraction of activators *p*^+^ = 0.5

In our investigation of the effects of modular network structure on the distribution of heritability, we simulated 10,000 PPM GRNs with generating parameters as given below:

- Number of genes *n* = 5000
- Number of groups *k ∼* Uniform(2, 100)
- Modularity term *m ∼* Uniform(1/*k*, 1)
- Expected regulators per gene *r ∼* Uniform(2, 10)
- Strength of regulation *γ ∼* Uniform(0.2, 0.5)
- Expected fraction of activators *p*^+^ *∼* Uniform(0, 1)

### 4.4 Modular directed acyclic scale-free graph

We use a modular scale-free graph generating algorithm to produce the synthetic DAGs for our full GRN model. This algorithm is an extension of our previous work, which we used to study experimental perturbations; the prior work is itself an extension of a directed scale-free network generating algorithm due to Bollobas [29, 30]. The algorithm we use here has an out-degree uniformity parameter *d*, and otherwise has parameters which are identical in interpretation to our re-parameterized PPM.

The output of this algorithm is a DAG with *n* nodes assigned into *k* groups. As with the PPM, the groups account for *m* percent of the edges in the graph, and the typical node has *r* neighbors — and further, the out-degree dispersion parameter *d* controls the spread of the out-degree distribution (**Fig. S7**). When *d* is near zero, the GRN has a concentrated regulatory architecture with hubs of outgoing regulation; when *d* is large, regulatory activity is spread more uniformly throughout the network, and there are many more regulators (but typically fewer hubs). Pseudocode for the algorithm is in **Appendix B**.

To specify a full GRN model using a DAG produced from this algorithm, and to produce gene expression from it, we follow the same procedure as used for the PPM DAGs. In our investigation of the joint effects of modular network structure and regulatory hubs, we simulated 10,000 GRNs using the following scheme to sample parameters:

- Number of genes *n* = 5000
- Degree uniformity log *d ∼* Uniform(log(1), log(30))
- Number of groups *k ∼* Uniform(2, 100)
- Modularity term *m ∼* Uniform(1/*k*, 1)
- Expected regulators per gene *r ∼* Uniform(2, 10)
- Strength of regulation *γ ∼* Uniform(0.2, 0.5)
- Expected fraction of activators *p*^+^ *∼* Uniform(0, 1)

We used these same 10,000 GRNs when matching the cumulative distribution of 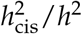 to that observed in real data [7].

### 4.5 Real data comparison

To assess the similarity of simulated GRNs to experimental data, we compared the expected cumulative distribution of *cis*-acting expression variance (*V*_cis_/*V*_G_) in each synthetic GRN to the cumulative distribution of the fraction of estimated 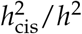 for 5,902 genes with a *cis*-eQTL from whole blood expression data [7]. We computed Kolmogorov-Smirnov (K-S) test statistics as a non-parametric measure of distance between the distributions from the simulated GRNs and that of real data. To determine which features of the gene expression model and network generating parameters were associated with closer matches to data, we further examined the GRNs at the 2.5% tail of the distribution of K-S test statistics (i.e., 250 lowest values) against the remaining 97.5% of simulated GRNs (from both PPM and scale-free GRNs; **Fig. 5, S11**).

### 4.6 Expression variance decomposition in synthetic networks

To further assess the genetic architecture of gene expression in example simulated GRNs, we used the same variance decomposition as in equation 14 — as a measure of the *trans*-effect *B*_*ji*_ of regulator *j* on gene *i*, we used the off-diagonal entries of the matrix *L*:

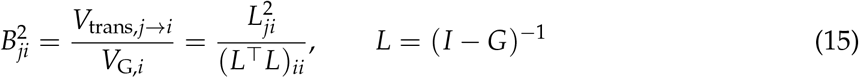

The leading *trans*-effect for gene *i* (as in **Figs. 5B** and **6E,F**) is the maximum effect over all regulators 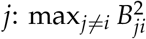. To assess the contributions of *trans*-regulators at varied distances *d* from gene *i* in the network, we compute the all-pairs shortest path distances between all genes in the network, then sum contributions from regulators at distance *d* — if *d*_*ji*_ is the distance from gene *j* to gene *i*, then the fraction of variance due to regulators at distance *d* is:

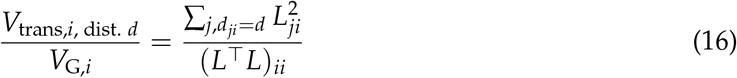

as is shown in **Figs. 6**, **S14**, **S15**.

## Appendix A Gene expression model

We model gene expression with a linear structural equation model (SEM) given a set of causal regulatory relationships specified by a directed acyclic graph (DAG) with *n* genes. We assume that the structure of the DAG is given and the parameters of the SEM are fully specified, and consider relaxations of this assumption later.

To start, consider the expression of a single gene *i*: suppose the gene harbors *q*_*i*_ independent cis-eQTLs (*q*_*i*_ *>* 0) and is affected by *r*_*i*_ regulators (*r*_*i*_ *≥* 0), which may or may not be independent. If we measure the expression *y*_*i*_ of this gene in a randomly chosen individual from population, we can write that

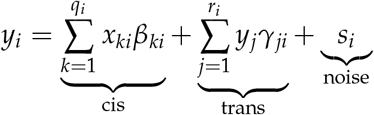

where above, *β*_*ki*_ and *x*_*ki*_ are respectively the effect of the *k*’th eQTL for gene *i* and an indicator variable for its genotype; *γ*_*ji*_ and *y*_*j*_ are the effect of the *j*’th regulator of gene *i* and its expression; and *s*_*i*_ is non-genetic noise with zero mean and variance *σ*^2^, which we assume is uncorrelated across genes.

Throughout this work, we assume that (1) each regulator acts consistently as an activator or repressor and has the same magnitude of effect (i.e., *γ*_*ji*_ = *p*_*j*_*γ*, with *p*_*j*_ denoting the sign for gene *j*), and that (2) the variance due to *cis*-acting eQTLs and transcriptional noise are the same for all genes (respectively 1, and *σ*^2^), and that (3) all genotypes and noise terms are independent. The variance for gene *i* is then

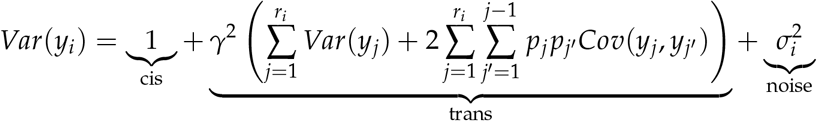

Now consider the expression of all genes in the entire network: if *G* is the weighted adjacency matrix of the provided DAG for the GRN, with entries *γ*_*ji*_ where regulatory relationships exist and zero otherwise, then we can more compactly write

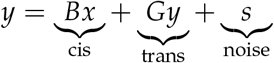

where *B* is the matrix of genetic effects from *cis*-eQTLs and *x* is a vector of their genotypes. We note that the above expression is valid since *G* is the matrix representation for DAG — this specifies a coherent set of equations for the expression, *y*_*i*_, of each gene, and guarantees that the matrix (*I − G*) is invertible. We can rearrange terms in the above to compute the variance of *y*:

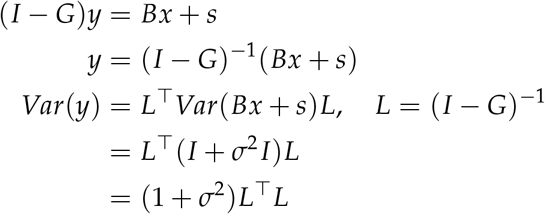

in which the marginal genetic variance of gene *i* can be read off the diagonal of *L*^*⊤*^*L*: *V*_*G*_(*y*_*i*_) = (*L*^*⊤*^*L*)_*ii*_. Moreover, this matrix product implies a decomposition of variance across genes in the network: since 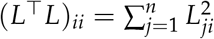 then the contribution of each gene *j* to the variance of gene *i* is

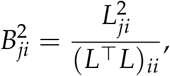

and the *cis*-acting fraction of genetic variance is correspondingly

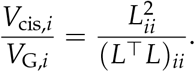

Furthermore, the equations above show that the genetic and non-genetic covariances have similar forms:

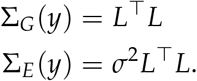

This means that any non-zero genetic contribution to variance has a corresponding non-genetic term which is scaled by a factor of *σ*^2^, and so the heritability for all genes in the network is constant: 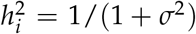. This parity also means that decompositions of genetic variance across genes in the network will generally correspond to decompositions of heritability, as long as this scaling factor is consistent across genes; for clarity in the main text, however, we refer to our work with genetic variance as such when using simulations from this model.

Furthermore, the matrix *L* = (*I* − *G*)^*−*1^ in the above expression is a total effects matrix for the structural equation model, which gives the marginal effect of the expression of gene *j* on the expression of gene *i*. More concretely, this equation corresponds to the Taylor expansion

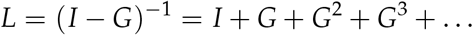

which enumerates all paths through the graph — moreover, since *G* is the adjacency matrix of a DAG, this is convergent since *G* is nilpotent (in particular, *G*^*n*^ is the zero matrix since *G* cannot have a path of length *n*). We use the squared entries of this matrix as the basis of our proxy for scaled *trans*-eQTL effect sizes. If *trans*-eQTL *k* for gene *i* has *cis*-effect *β*_*kj*_ on gene *j*, then the squared *trans*-effect is 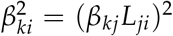. In practice, however, these effects are computed relative to the variance in expression of the focal gene (*j* for the *cis* effect, or *i* for the *trans* effect). Hence, to compare the relative scales of *cis* and *trans* effect sizes in this model, we use the proportion of variance of gene *i* explained by gene *j*,

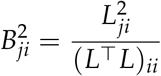

as a proxy for *trans* effects on gene *i* from *cis*-eQTLs of gene *j*. This corresponds to the assumption that *cis* effects *β*_*kj*_, *β*_*k*_*′* _*i*_ (of eQTL *k* on gene *j*, and eQTL *k′* on gene *i*) are equal.

## Appendix B Modular scale-free directed acyclic graphs

### Algorithm 1

Modular scale-free DAG

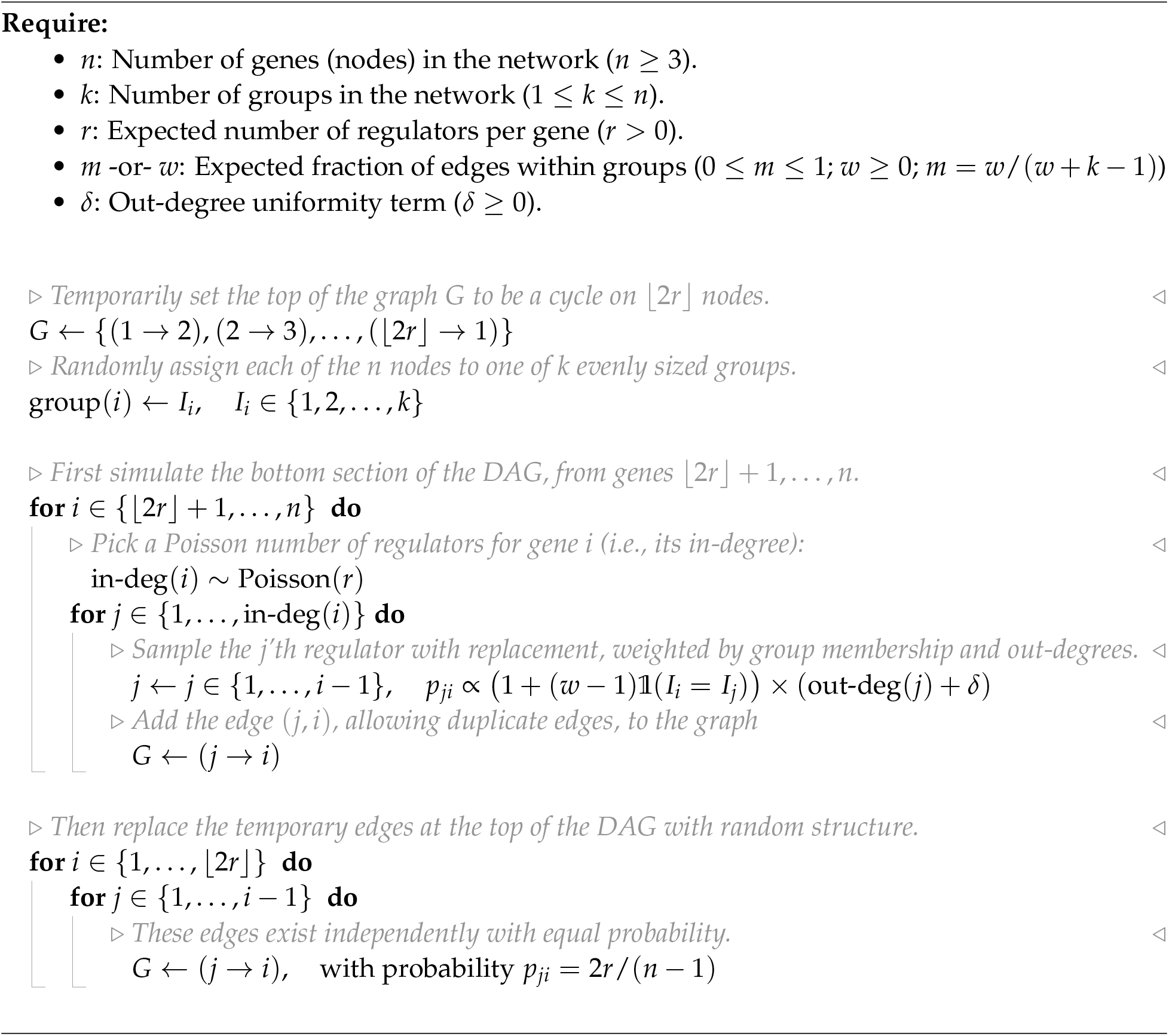

## 5 Supplementary Information

**Figure S1:**
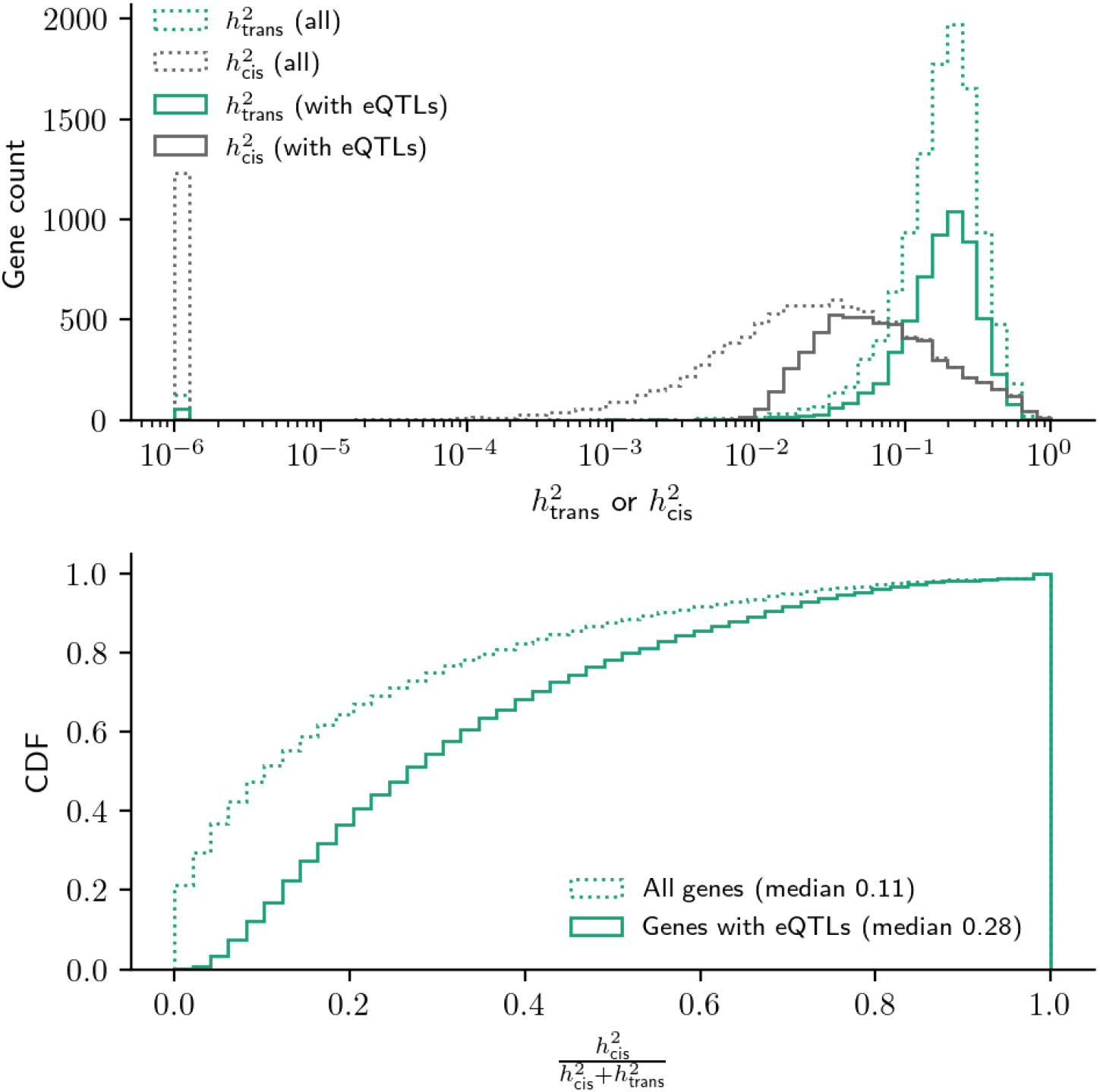
Distribution of *cis* and *trans* heritability for genes with and without eQTLs. Data from (Ouwens *et. al*., 2020). **(A)** Distribution of 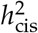 and 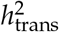 for all 11,353 genes with available data from the study (dotted lines) or for all 5,902 genes in the analysis subset from **Fig. 1**, with a detected eQTL (solid lines). **(B)** Distribution of *cis*-heritability fraction from these same gene sets.

**Figure S2:**
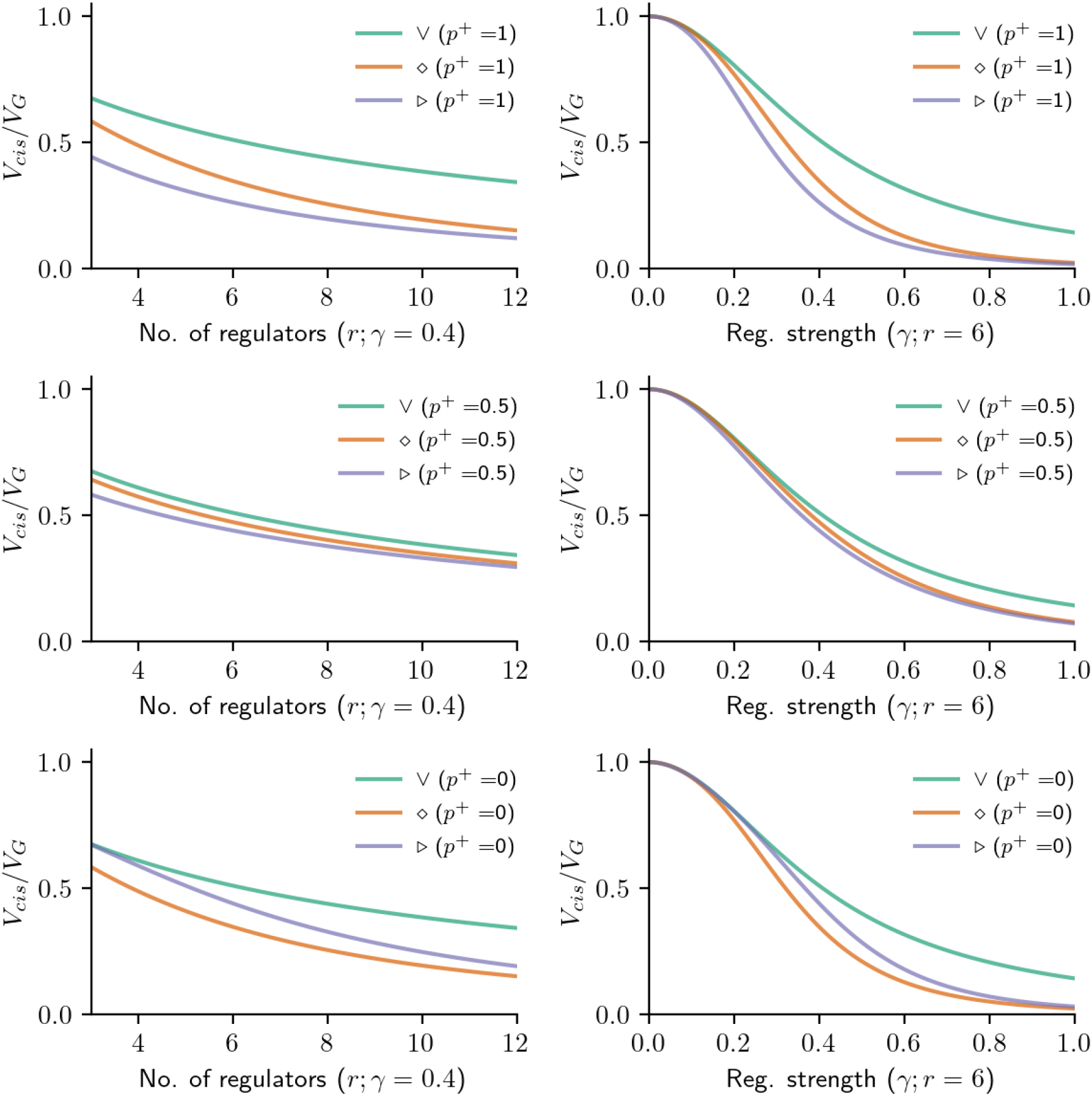
Effect of motifs with different fractions of activators. Representative effects of the number of regulators (*r*; left panels) and the strength of regulation (*γ*; right panels) on the distribution of *cis*-acting variance for the three motifs in **Fig. 2**. The underlying mathematical equations for vee, triangle, and diamond motifs are as in the main text and corresponding figure — here, the expressions are also stratified by representative fractions of activators *p*^+^ (the panels in the top row are exactly as in **Fig. 2**).

**Figure S3:**
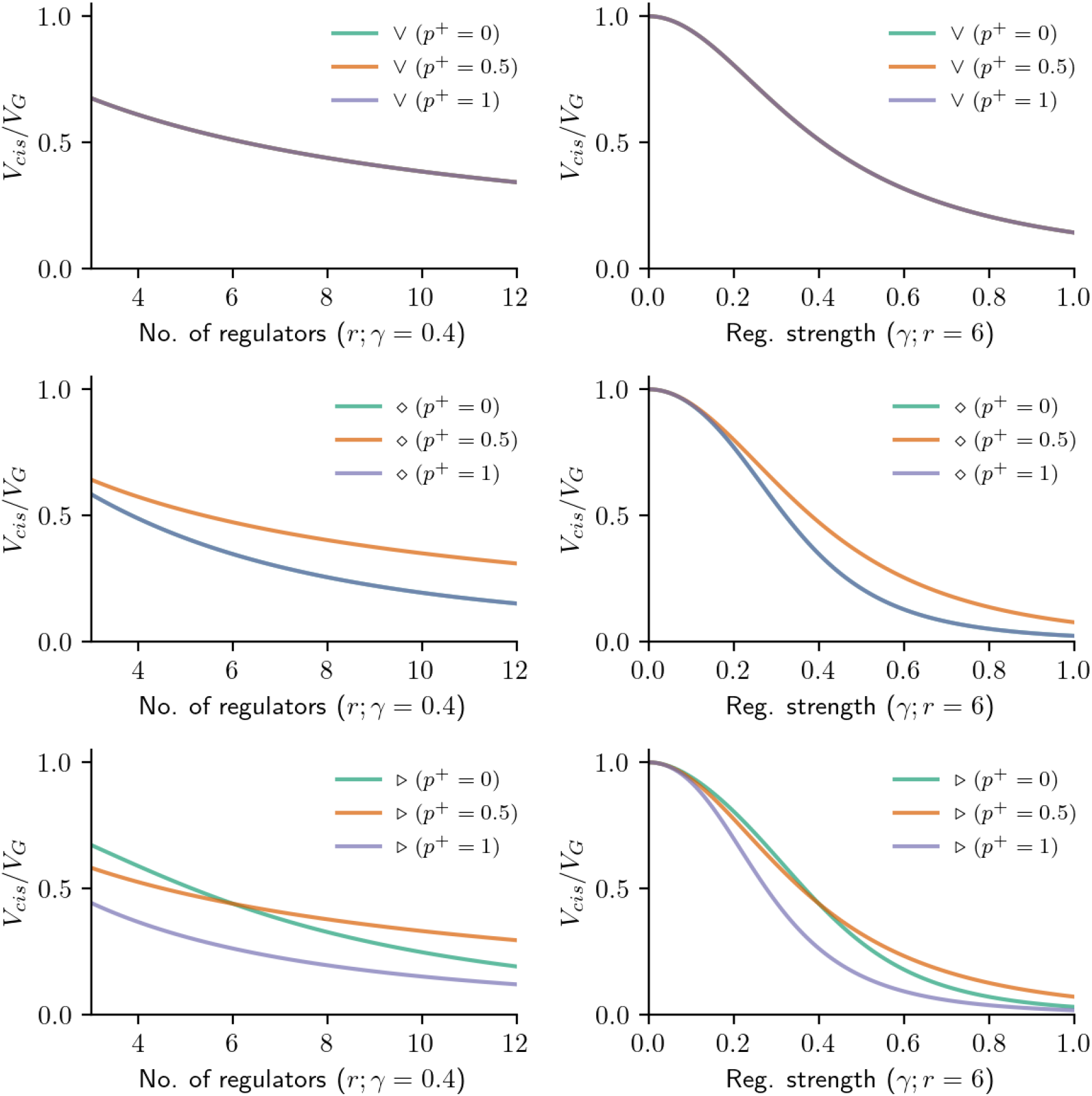
Effect of regulatory sign on different motifs. Representative effects of the number of regulators (*r*; left panels) and the strength of regulation (*γ*; right panels) on the distribution of *cis*-acting variance for the three motifs in **Fig. 2**. The lines in this plot are the same as in **Fig. S2**, but subpanels here correspond to the separate motifs (vee, diamond, and triangle, in each row) rather than distinct values for the fraction of activators (*p*^+^).

**Figure S4:**
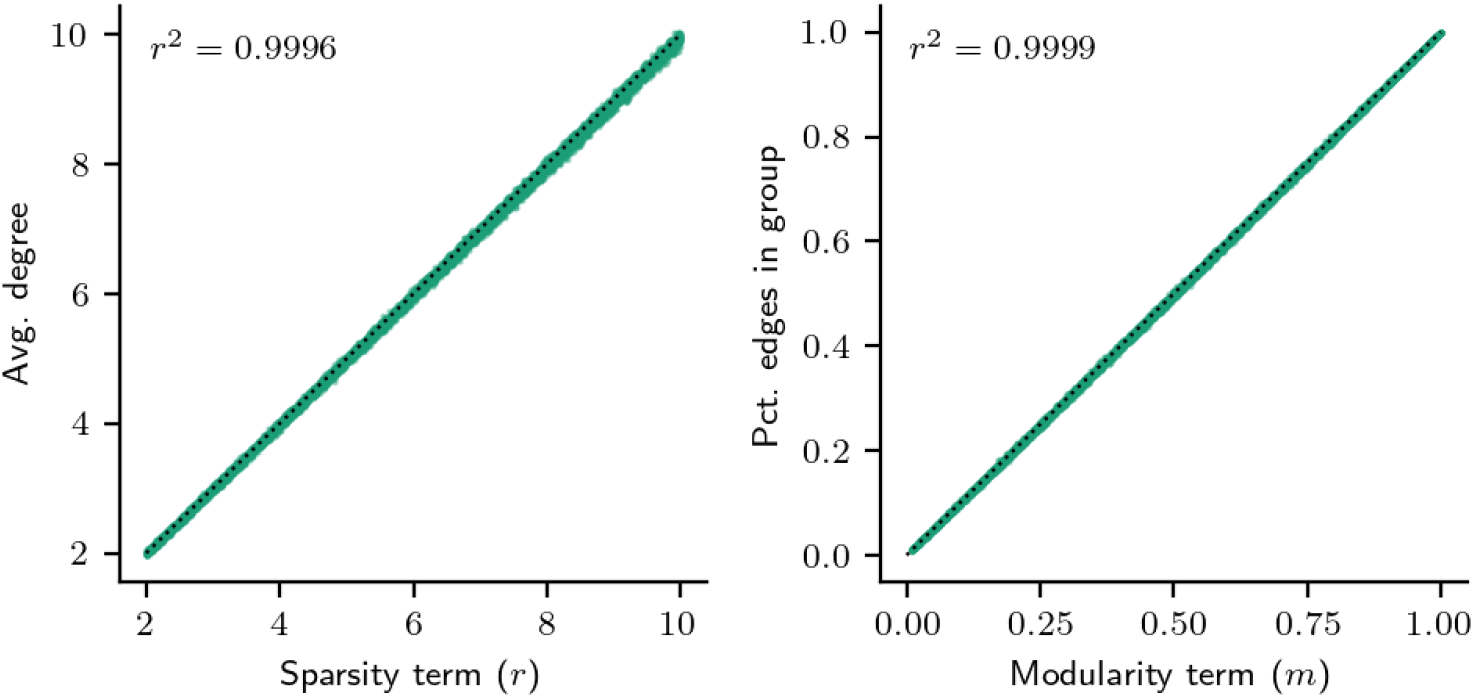
Parameters of the planted partition model (PPM) control key graph properties. Values for the sparsity term (*r*; left panel) and modularity term (*m*; right panel) in the 10,000 synthetic GRNs generated using the planted partition model (see **Methods**). The sparsity term *r* is extremely correlated with the average degree in the resulting GRN, and the modularity term *m* is extremely correlated with the resulting fraction of edges in the GRN that are drawn between genes in the same group (rather than genes in separate groups).

**Figure S5:**
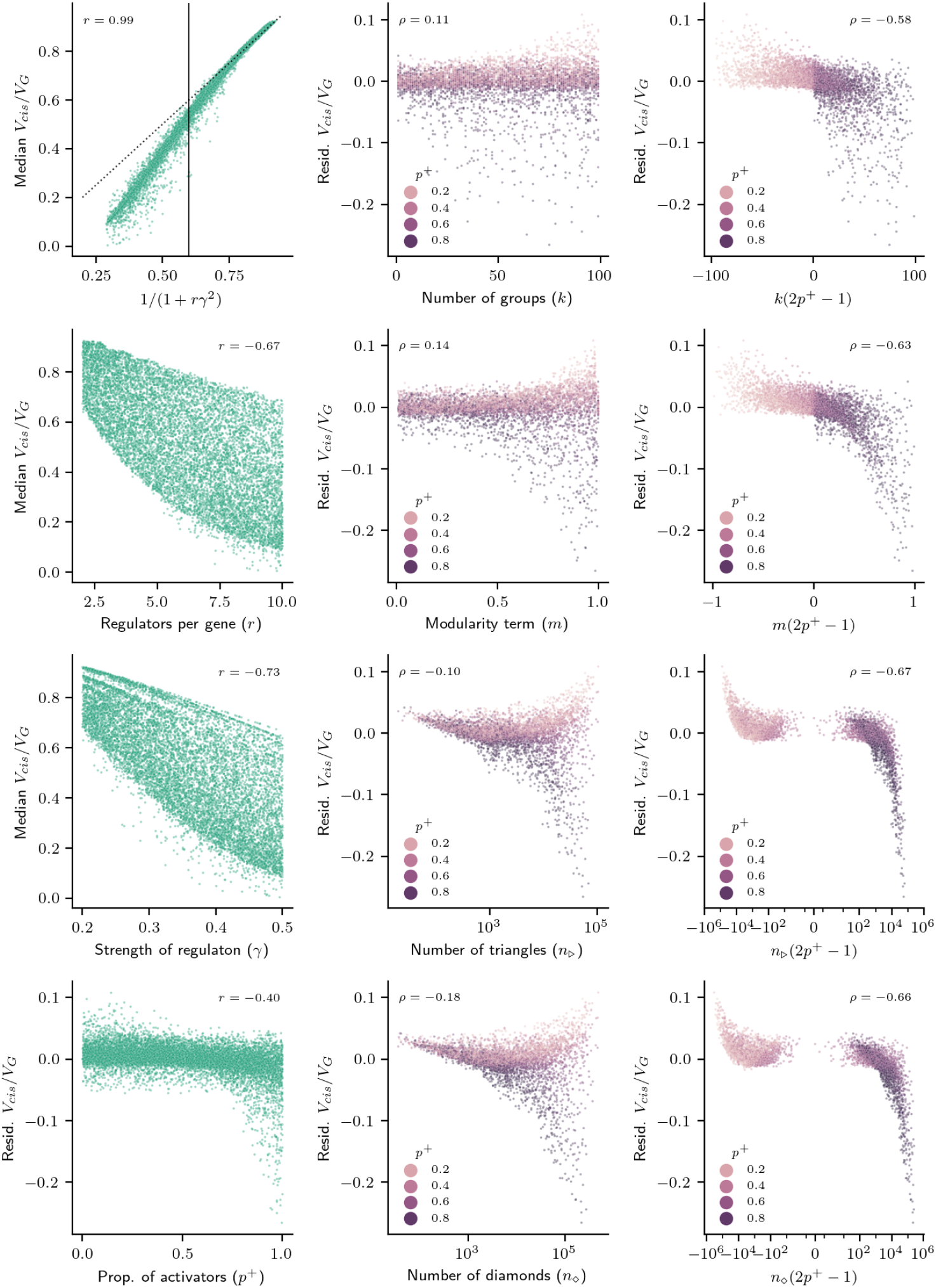
Parameters of the planted partition model (PPM) affect the distribution of heritability. Relationships between properties of GRNs produced using the PPM and summaries of the median fraction of *cis*-acting expression variance in the corresponding 10,000 GRNs. Subpanels are annotated by the Pearson (*r*) or Spearman (*ρ*) correlation between the quantities shown on the axes — median *V*_cis_/*V*_G_ denotes the untransformed value, while “Resid. *V*_cis_/*V*_G_” denotes the residual having regressed out directed effects as 1/(1 + *rγ*^2^). Networks in the middle and right columns are shown stratified by the fraction of activators *p*^+^ and are subsetted to GRNs with lower expected contributions from direct effects (1/(1 + *rγ*^2^) *<* 0.6, denoted by a solid vertical line in the top left panel (the dashed line is *y* = *x*); these are also the networks shown in **Fig. 3**.

**Figure S6:**
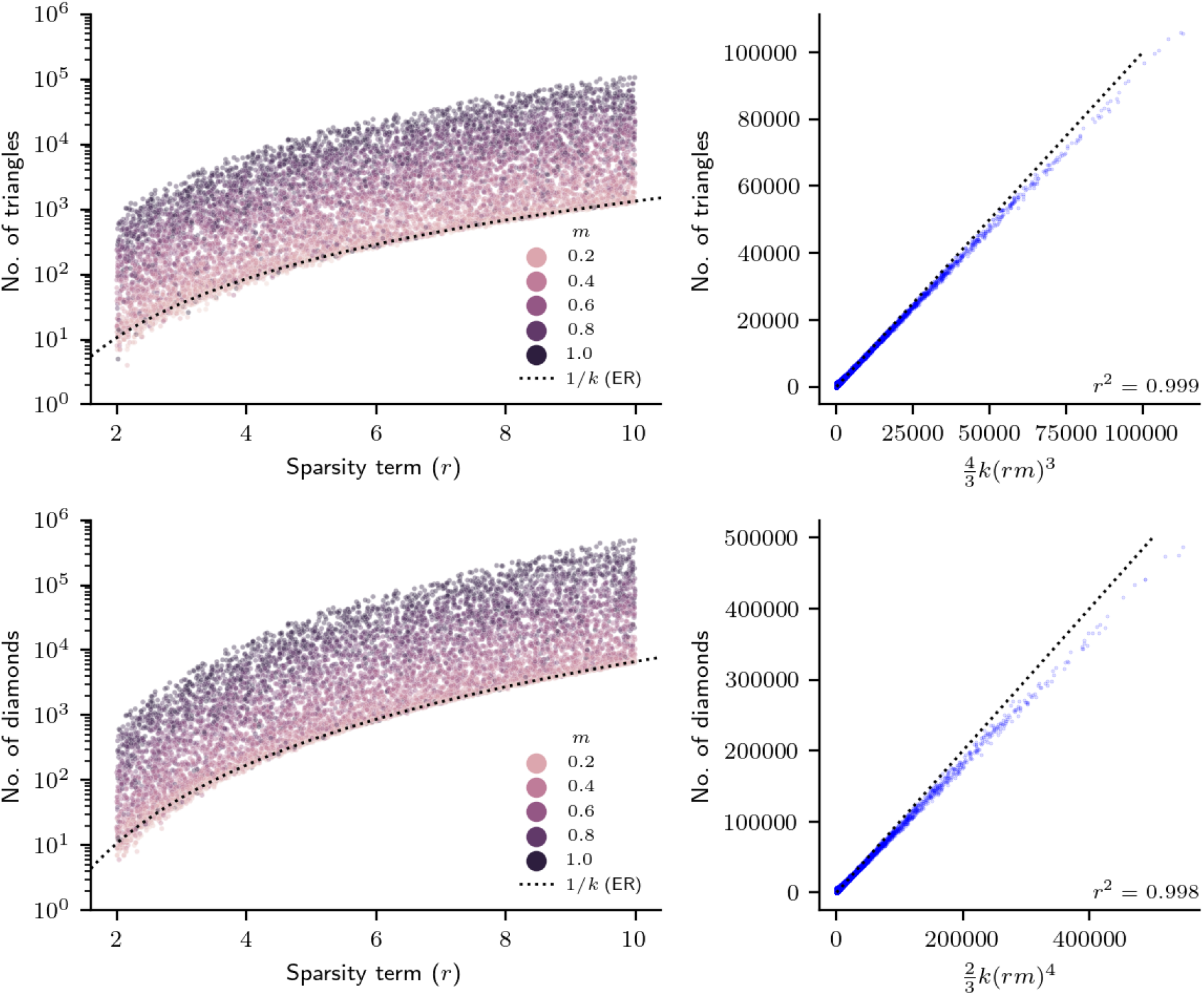
Parameters of the planted partition model (PPM) affect the number of motifs in the network. Relationships between the sparsity (*r*) and modularity (*m*) parameters of the PPM and the resulting number of triangle and diamond motifs (defined as diagrammed in **Fig. 2**) in 10,000 PPM GRNs. In both plots, each point is a network — the left panels show the interaction between *r* and *m* (note that *m* = 1/*k* is the minimum value simulated in the study, and corresponds to the binomial graph); the right panels show the relationship between motif counts and the leading term of a mathematical approximation to the expected number of motifs.

**Figure S7:**
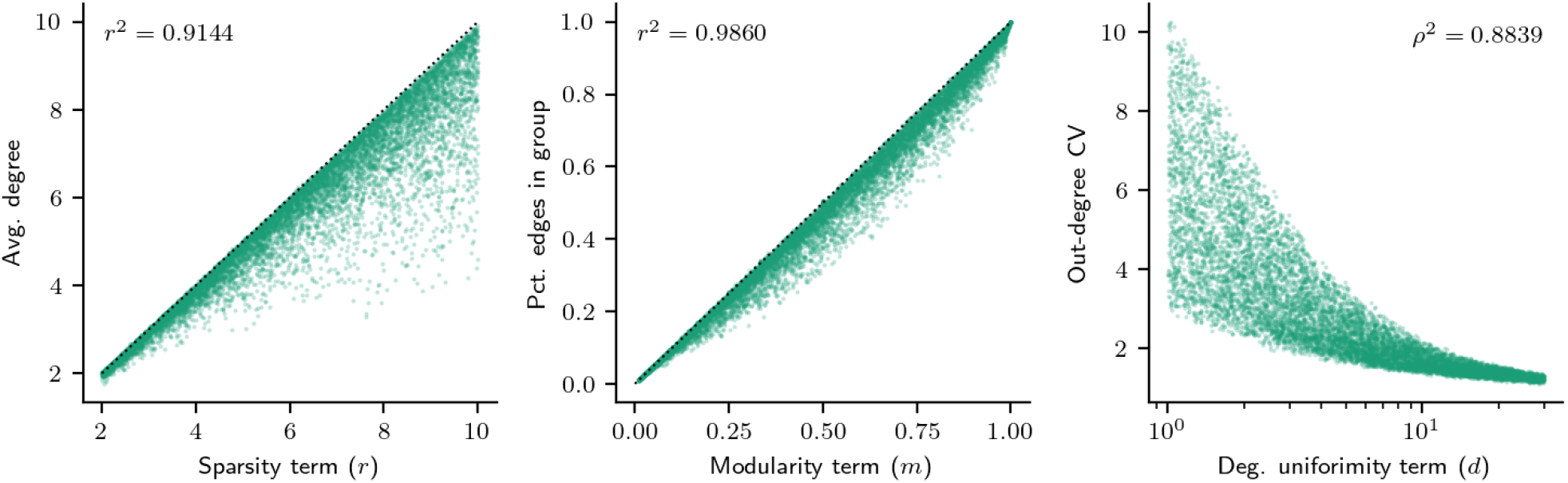
Parameters of the modular scale-free graph model control key graph properties. Values for the sparsity term (*r*; left panel) and modularity term (*m*; right panel) in the 10,000 synthetic GRNs generated using the modular scale-free graph generating algorithm (see **Methods**). The sparsity term *r* is extremely correlated with the average degree in the resulting GRN. The modularity term *m* is extremely correlated with the resulting fraction of edges in the GRN that are drawn between genes in the same group (rather than genes in separate groups). The degree uniformity term *d* is strongly correlated with the coefficient of variation (CV) of the out-degree distribution — here, CV is the standard deviation of the degree distribution over its mean, and a larger CV corresponds to a more dispersed distribution.

**Figure S8:**
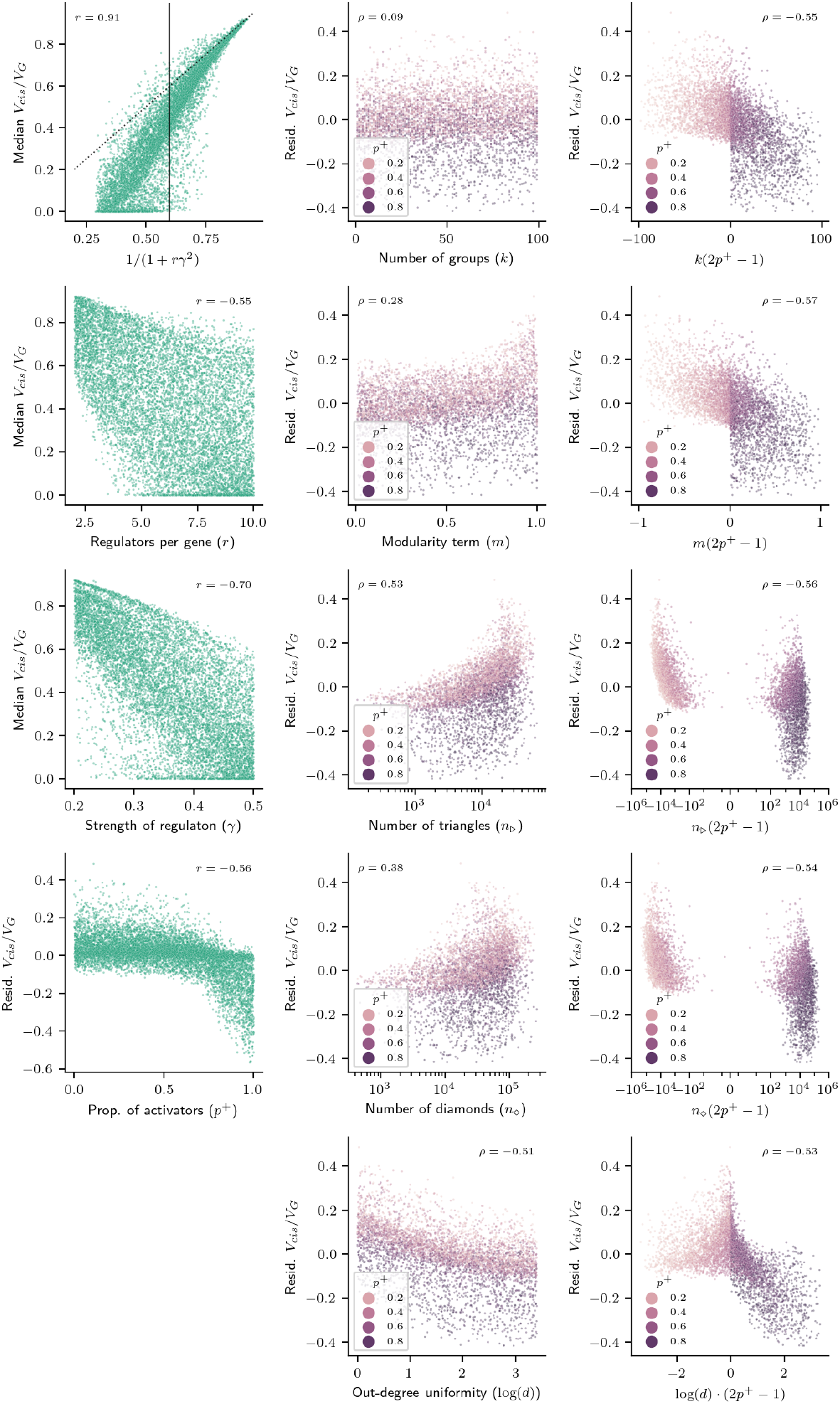
Parameters of the modular scale-free graph model affect the distribution of heritability. Relationships between properties of GRNs produced using the modular scale-free graph model and summaries of the median fraction of *cis*-acting expression variance in the corresponding 10,000 GRNs. Subpanels are annotated by the Pearson (*r*) or Spearman (*ρ*) correlation between the quantities shown on the axes — median *V*_cis_/*V*_G_ denotes the untransformed value, while “Resid. *V*_cis_/*V*_G_” denotes the residual variance after regressing out direct effects as 1/(1 + *rγ*^2^). Networks in the middle and right columns are shown stratified by the fraction of activators *p*^+^ and are subset to GRNs with lower expected contributions from direct effects (1/(1 + *rγ*^2^) *<* 0.6, denoted by a solid vertical line in the top left panel (the dashed line is *y* = *x*); these are also the networks shown in **Fig. 4**.

**Figure S9:**
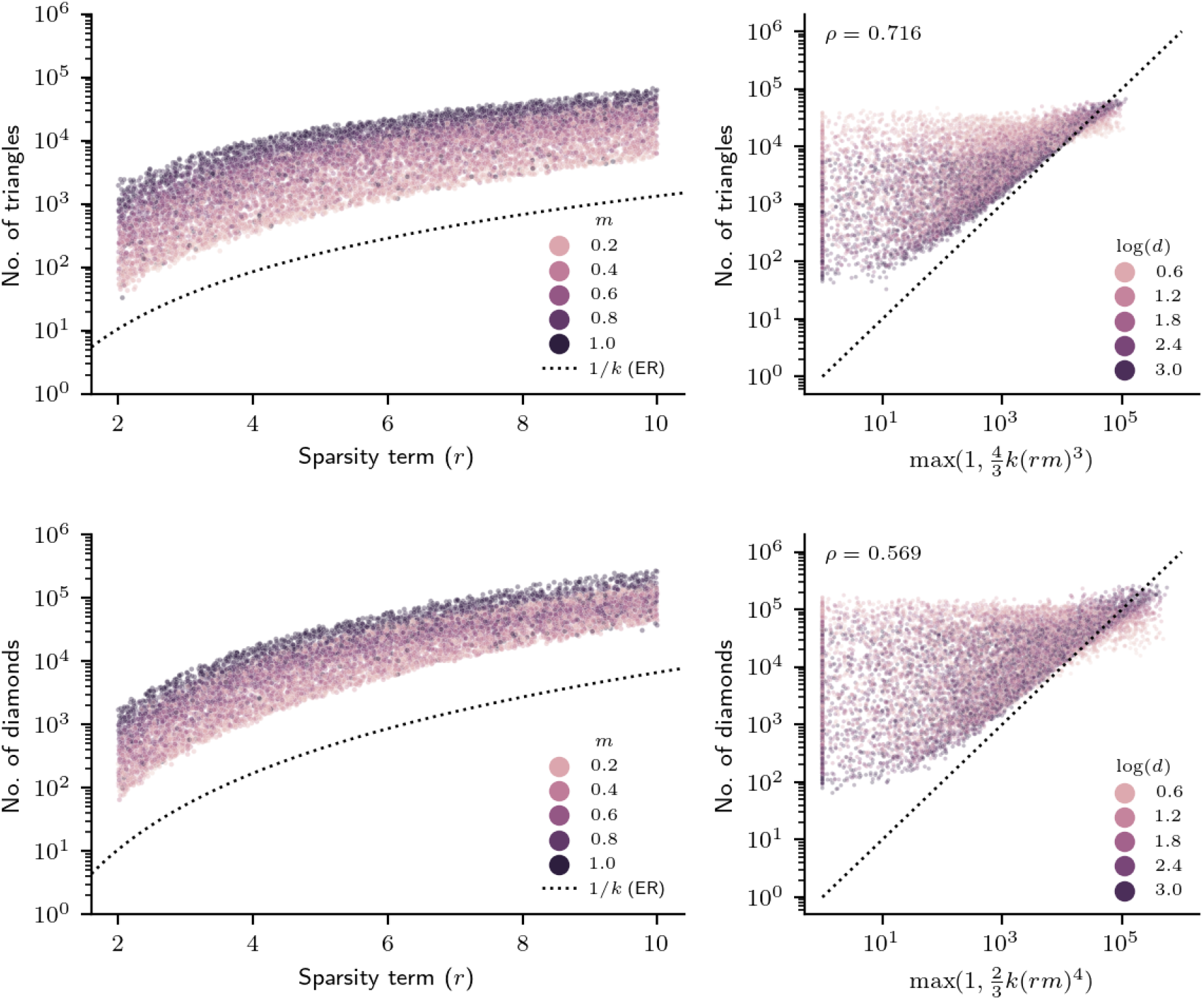
Parameters of the modular scale-free graph model affect the number of motifs in the network. Relationships between the sparsity (*r*) and modularity (*m*) parameters of the modular scale-free graph model and the resulting number of triangle and diamond motifs (defined as diagrammed in **Fig. 2**) in 10,000 GRNs. In both plots, each point is a network — the left panels show the interaction between *r* and *m* (note that *m* = 1/*k* is the minimum value simulated in the study, and corresponds to the value which yields the binomial graph in the PPM); the right panels show the relationship between motif counts and the leading term of a mathematical approximation to the expected number of motifs to the PPM, and that smaller values of *d* (which introduce hubs into the graph) drive deviations from this expected value.

**Figure S10:**
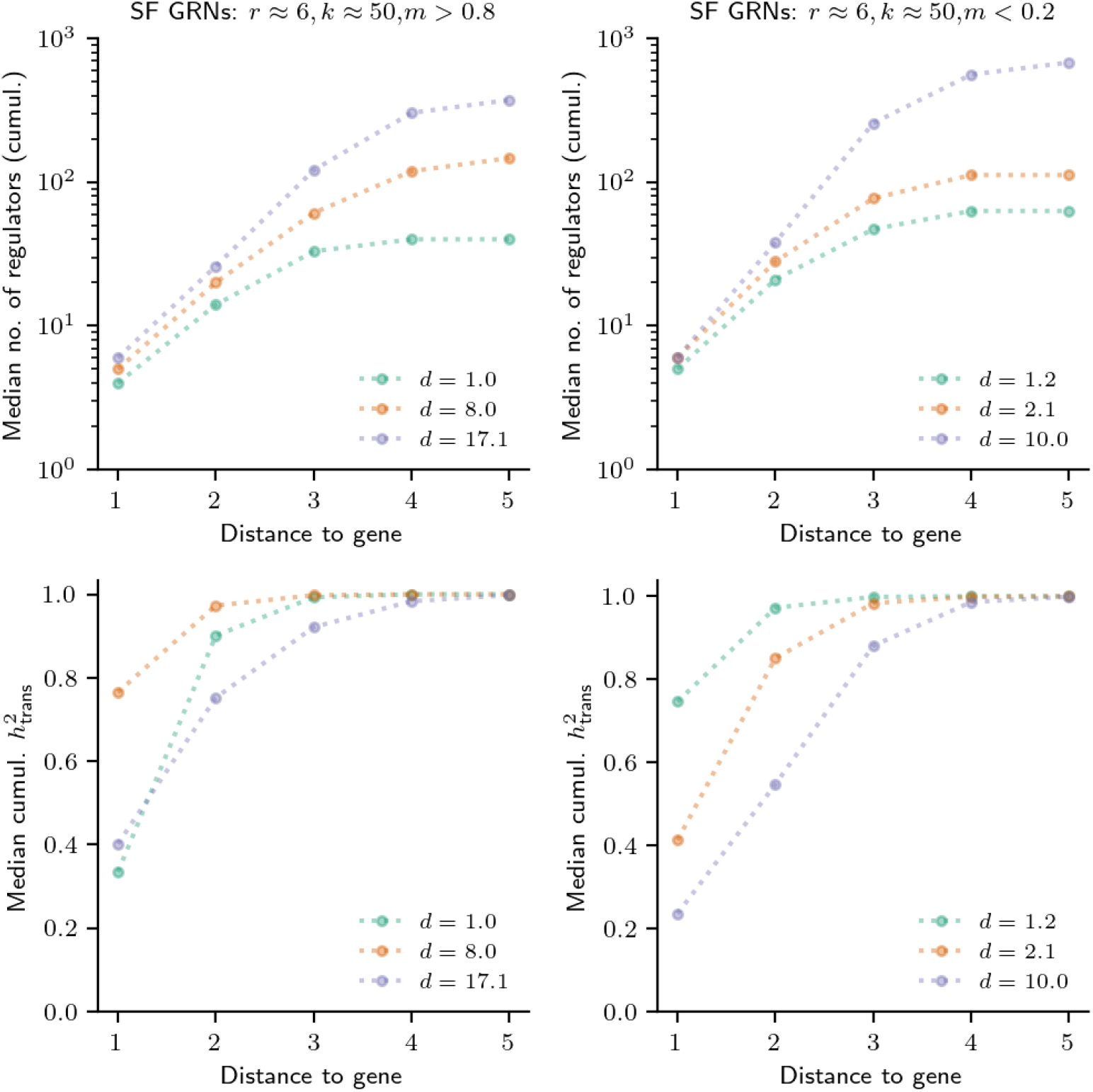
Degree uniformity alters the distribution of path lengths in the network. Median number of regulators (top panels) and cumulative *trans*-acting expression variance (bottom panels) in six example modular scale-free GRNs, matched to have similar sparsity (*r* ≈ 6) and group structure (*k* ≈ 50, *m >* 0.8 or *m <* 0.2), but vary in other regulatory terms (*d, γ, p*^+^). Medians are over genes in the middle of the topological sorted order of the DAG (indexes 2000 to 3000). GRNs with regulatory hubs (smaller values of *d*) have fewer distant *trans*-regulators and *trans*-acting heritability is thus typically closer to a given gene in the network.

**Figure S11:**
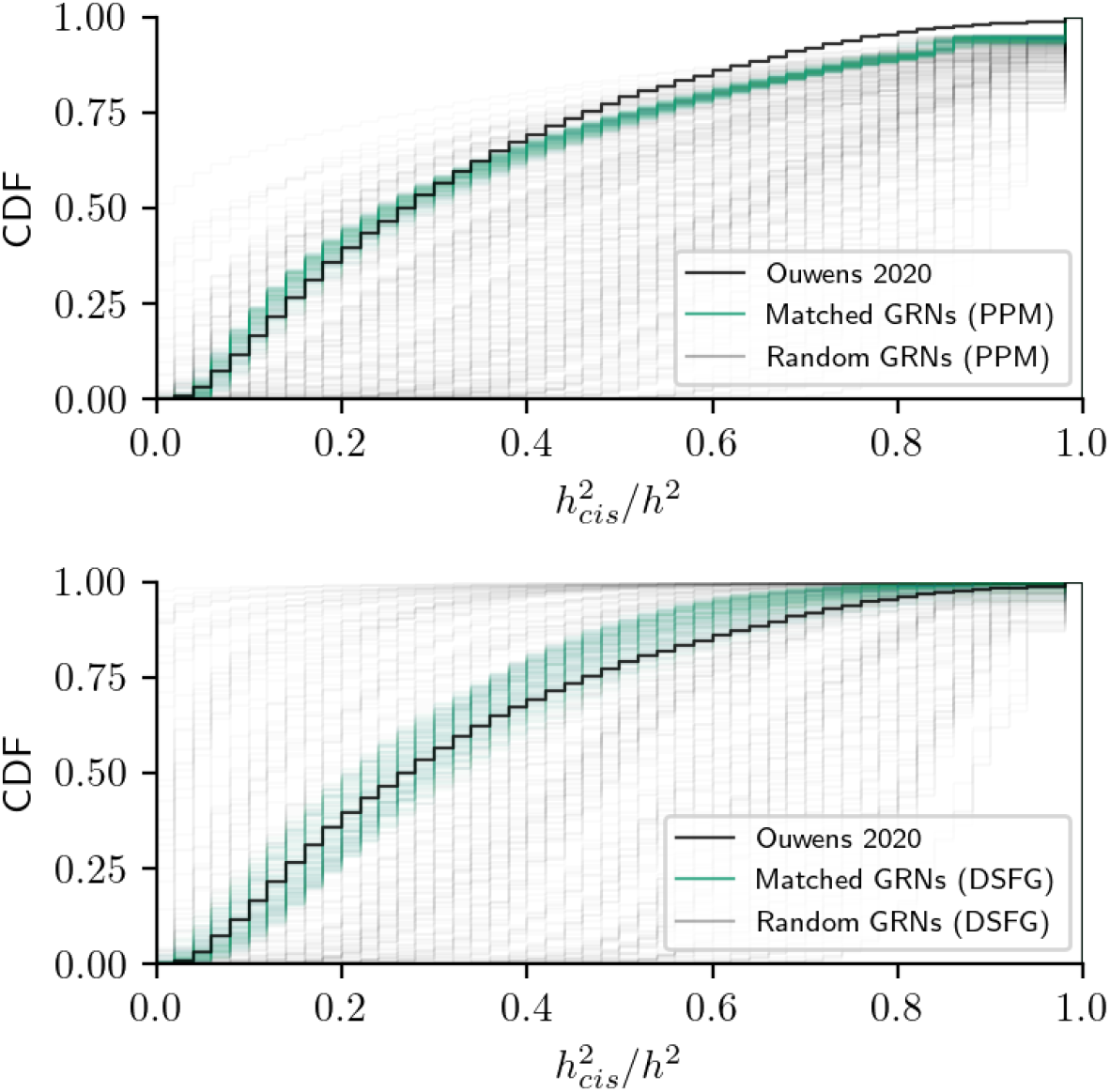
Distribution of heritability in synthetic GRNs and for whole-blood gene expression. Distribution of the fraction of *cis*-acting heritability 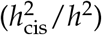 from real data (Ouwens *et. al*. 2020) and from synthetic GRNs generated using the planted partition model (PPM; top panel) or the directed scale-free graph generating algorithm (SF; bottom panel). In both plots, teal lines are the 250 GRNs closest to the distribution from data (lowest K-S test statistics; see **Methods**), and grey lines are a random sample of 250 GRNs from the other 9,750 simulated GRNs.

**Figure S12:**
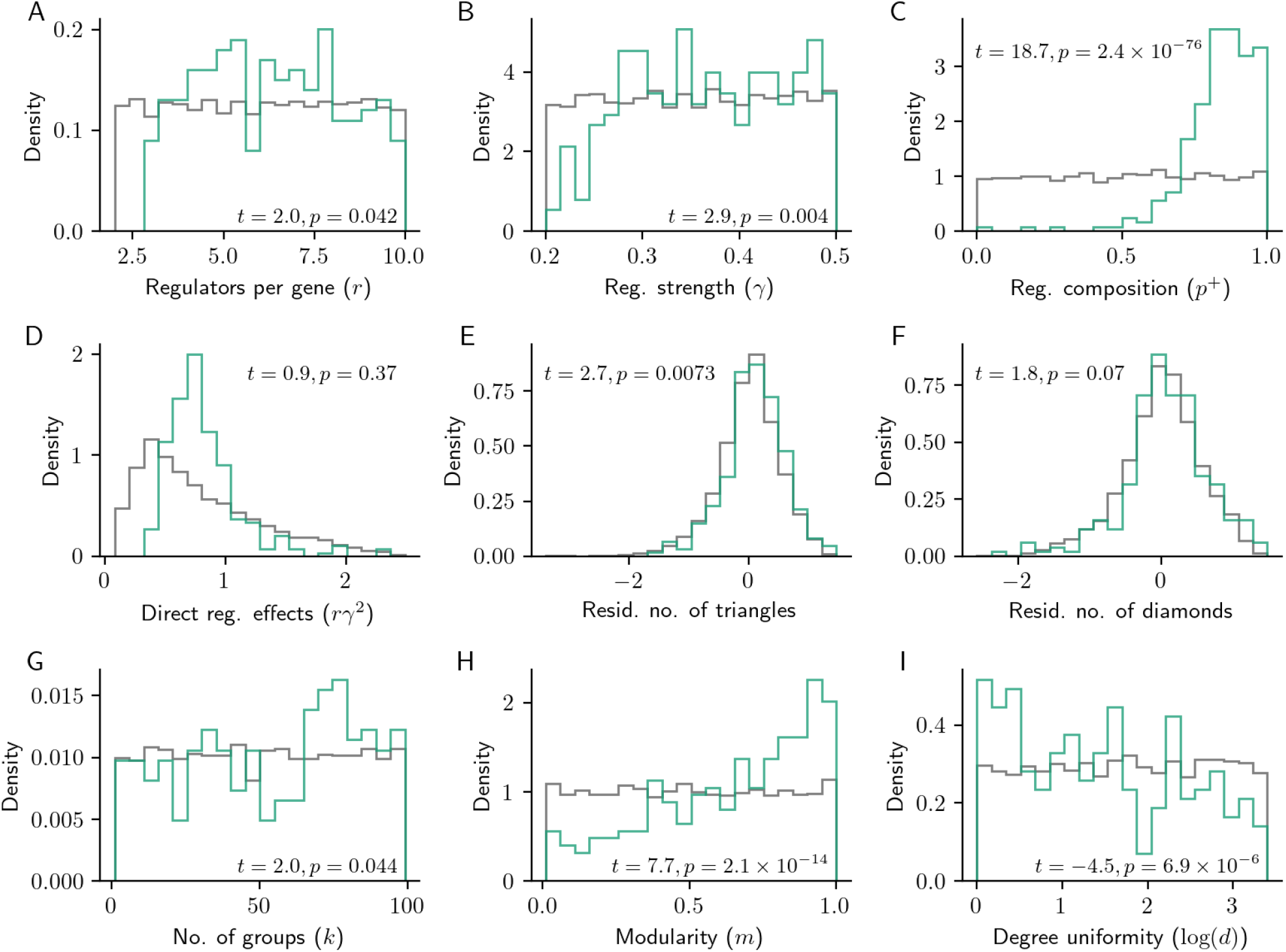
Properties of synthetic GRNs that best resemble real data. Additional properties of the 250 GRNs that were best matched to the observed distribution of *cis*-acting heritability fractions (shown in teal, as in **Fig. 5**). Each panel is annotated with results from a two-sample *t*-test for a difference in mean with the remaining 9,750 GRNs (shown in grey).

**Figure S13:**
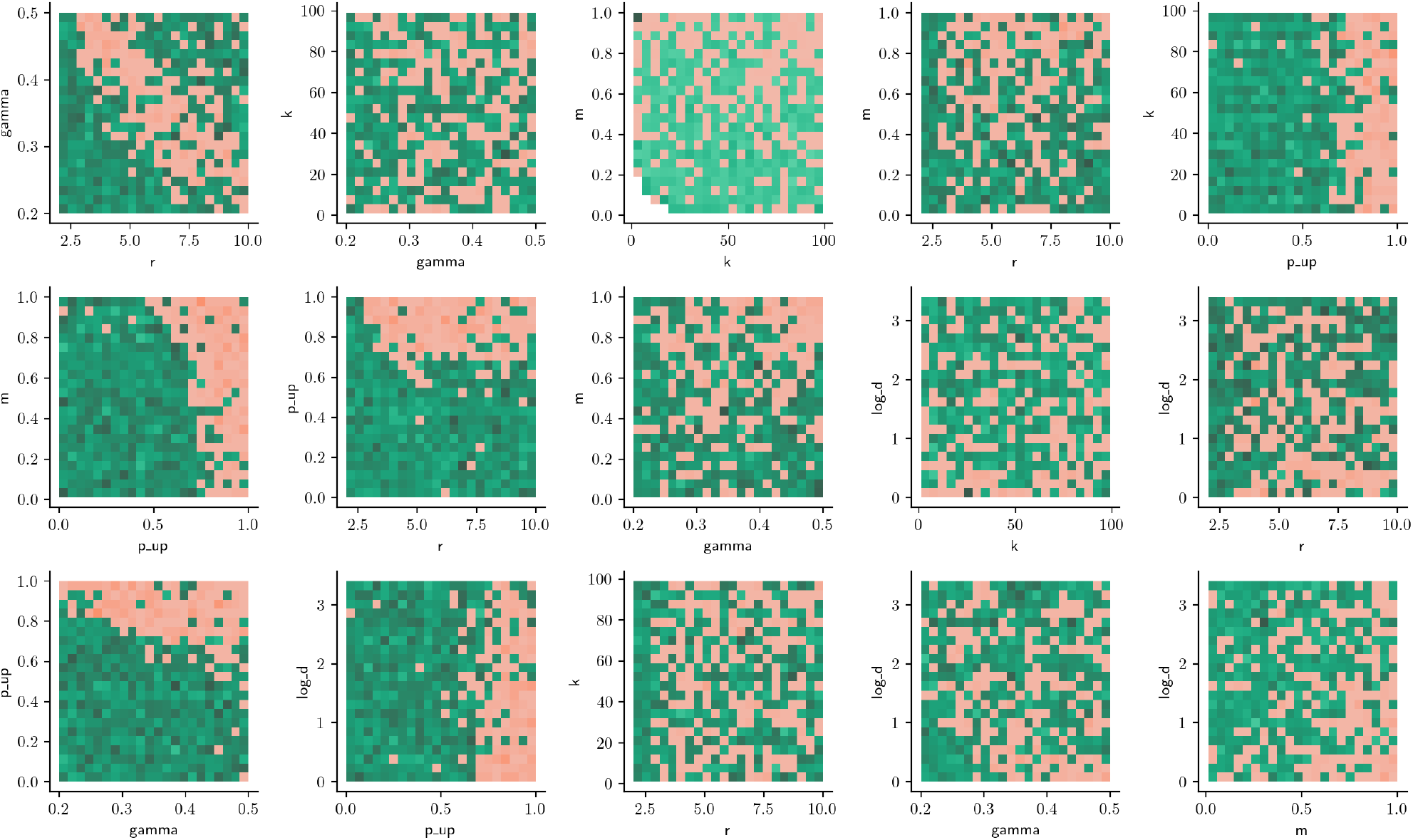
Interactions between properties of synthetic GRNs that best resemble real expression heritability data. Pairwise interactions between properties of the 250 GRNs that were best matched to the observed distribution of *cis*-acting heritability fractions. In each subpanel, red denotes an enrichment of matched GRNs in and green denotes depletion.

**Figure S14:**
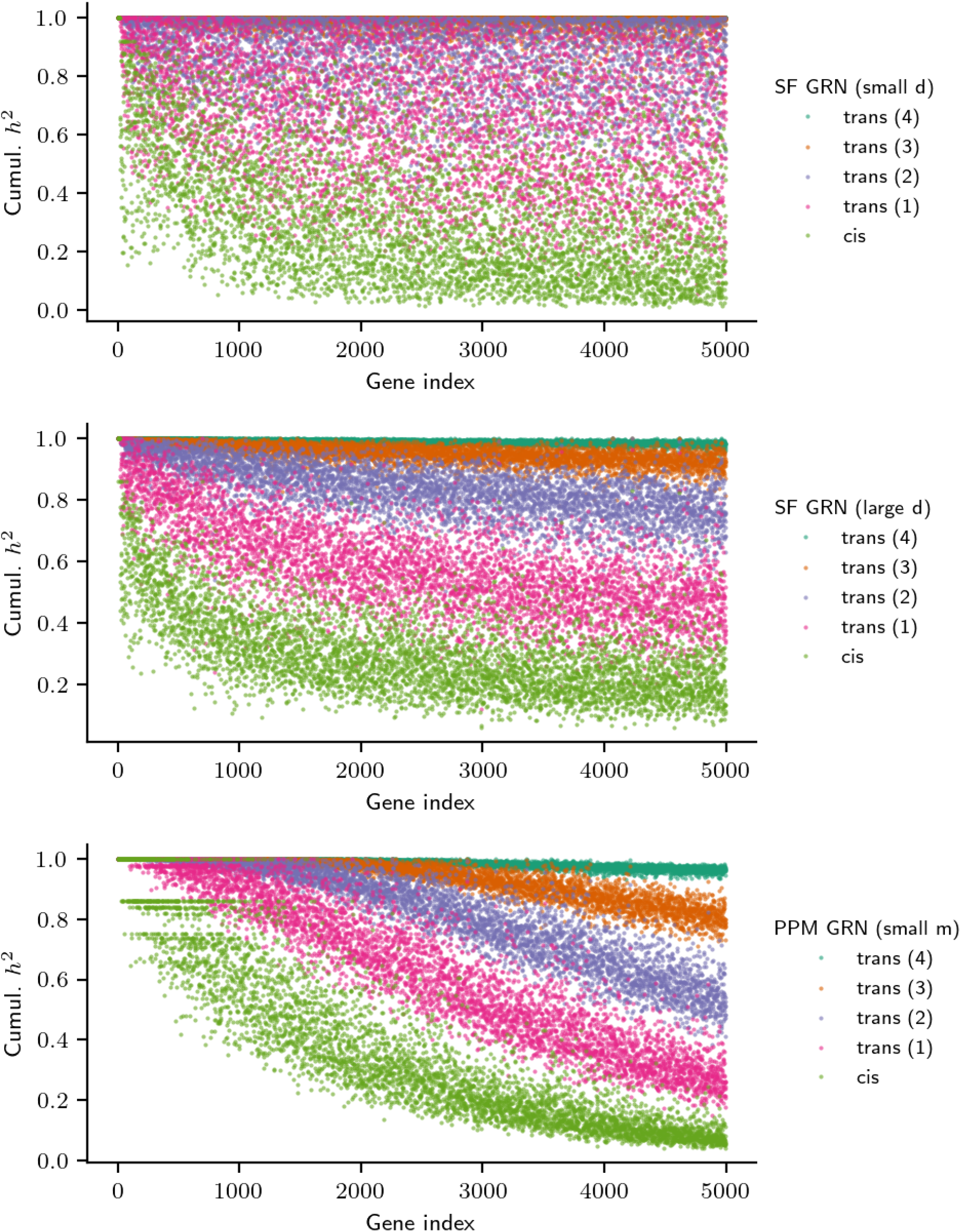
Gene-level heritability distribution over network distances in three example GRNs. Cumulative heritability in *cis* and in *trans*, stratified by distance, for all genes in the three example networks from **Fig. 6**.

**Figure S15:**
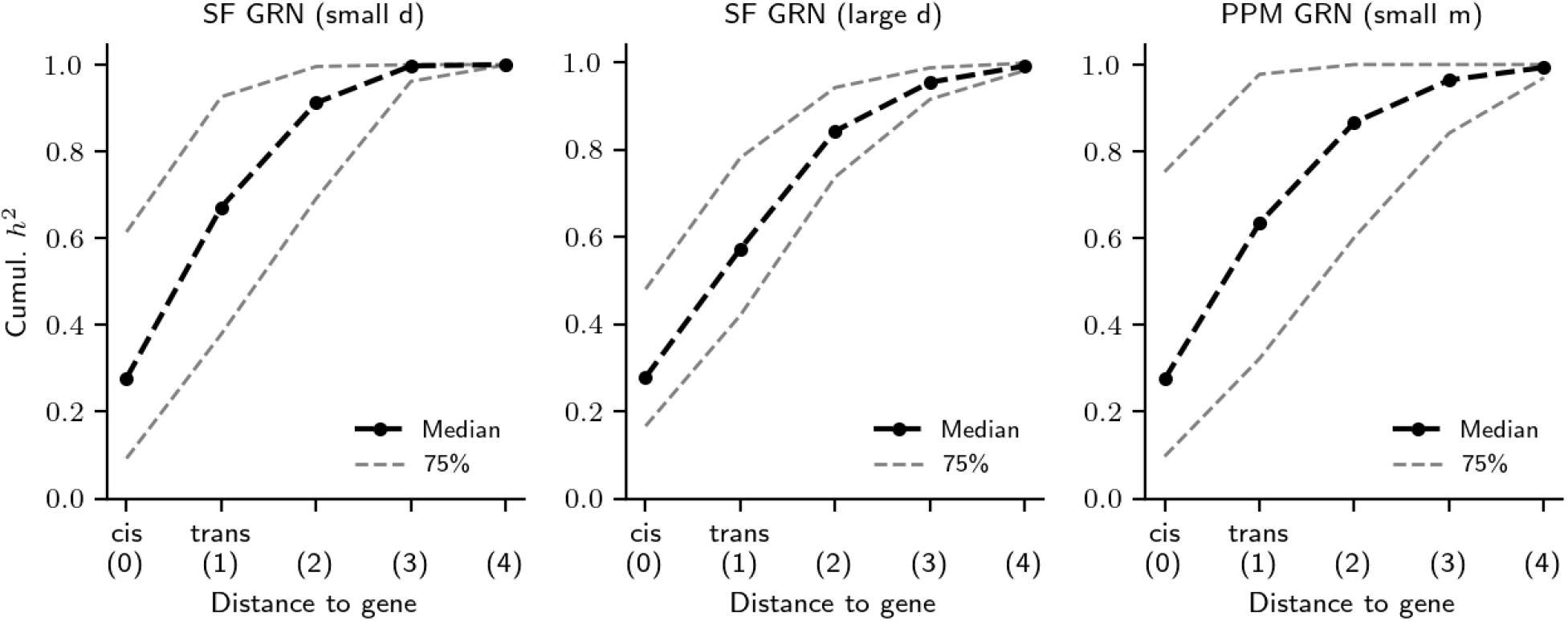
Cumulative heritability distribution over network distances in three example GRNs. Cumulative heritability in *cis* and in *trans*, stratified by distance, for the median gene (or quartiles) in the three example networks from **Fig. 6B** — note that the medians and quantiles are computed separately for each tick in the *x*-axis (i.e., the gene with the median *cis*-acting heritability fraction may not be the gene with the median one-hop *trans*-acting heritability fraction).

